# Complementary evidence from historical and contemporary gene dispersal reveals contrasting population dynamics in a tropical tree species

**DOI:** 10.64898/2026.03.23.713184

**Authors:** Julien Bonnier, Myriam Heuertz, Stéphane Traissac, Olivier Brunaux, Olivier Lepais, Valérie Troispoux, Emilie Chancerel, Zoé Compagnie, Niklas Tysklind

## Abstract

Gene flow shapes the demographic stability and evolutionary potential of tropical forest trees, yet its dynamics may differ depending on the temporal scale at which it is assessed. We combined spatial genetic structure (SGS), parentage analyses, and reproductive success metrics to investigate historical and contemporary gene dispersal in four populations of *Dicorynia guianensis* across French Guiana, encompassing sites differing in environment and management history. A total of 1,528 individuals were genotyped using 66 nuclear and 23 plastid microsatellite markers, enabling high-resolution inference of biparental and maternal gene dispersal. Historical mating and dispersal parameters inferred from SGS revealed marked contrasts among populations. Some populations exhibited high historical gene dispersal distances and weak spatial genetic structure, whereas others showed stronger SGS and long-term aggregative dispersal patterns. Contemporary parentage analyses further highlighted differences in seed and pollen dispersal distances, parent assignment rates, and reproductive skew. In certain populations, pronounced reproductive inequality and reduced effective connectivity were observed, while others displayed more balanced reproductive contributions. By jointly evaluating long-term dispersal legacies and present-day reproductive patterns, our study demonstrates the value of combining indirect and direct genetic approaches to assess population dynamics and conservation status in tropical forest trees. This multi-temporal perspective provides a comprehensive basis for long-term monitoring and sustainable management in heterogeneous tropical landscapes.

## 1. Introduction

Gene flow is a fundamental evolutionary process shaping the spatial distribution of genetic variation within and among plant populations. It promotes genetic connectivity by enabling the movement of alleles across landscapes, and counteracts the effects of genetic drift and local inbreeding (Petit & Hampe, 2006). Exchange of genetic material introduces novel alleles and supports the maintenance of genetic diversity. This is crucial for the adaptive potential and resilience of forest species, especially in changing environments (Lowe et al., 2005). In tropical forest ecosystems, gene flow helps buffer the effects of demographic fluctuations and local disturbances, ensuring population viability in the long term. The ability of tree populations to respond to environmental variation, including climate change, habitat degradation, and fragmentation, is related to their levels of gene flow and the spatial structure of their genetic diversity (Savolainen et al., 2007; Ward et al., 2005). From a management perspective, understanding gene flow is essential for designing strategies aimed at conserving genetic resources, enhancing connectivity among forest patches, and maintaining evolutionary potential in the face of ongoing anthropogenic pressures (Allendorf, 2017; Escudero et al., 2003).

The study of gene flow in plant populations has historically relied on indirect methods, particularly those based on patterns of genetic divergence. Classical population genetic models such as the island model or the isolation by distance model infer gene flow from the degree of genetic divergence between geographically isolated populations, or from the increase of population divergence with geographical distance (Rousset, 1997; Wright, 1943, 1965). These approaches assume drift-migration equilibrium and typically reflect historical patterns of gene flow integrated over many generations, which makes them useful for studying long-term evolutionary processes but limits their capacity to detect recent demographic changes or fine-scale reproductive dynamics. To understand more local processes, spatial genetic structure (SGS) analysis within populations has become a widely used indirect approach. SGS quantifies how genetic relatedness decays with geographic distance within a population, offering insights into how restricted dispersal and local demography shape genetic patterns over time (Epperson, 1992; Rousset, 2000; Vekemans & Hardy, 2004). In plants, SGS arises primarily from seed and pollen dispersal patterns but is also influenced by ecological conditions, demographic fluctuations, and life-history traits (Anderson et al., 2010; Hardy et al., 2006). SGS does not measure dispersal events directly but provides a powerful tool to infer historical effective gene dispersal, especially in cases where direct observation is difficult or unfeasible, such as in tropical trees (Dick et al., 2008).

Direct approaches such as parentage analysis and mating models have emerged as robust methods for characterizing contemporary dispersal events and gene flow. These approaches involve genotyping offspring and candidate parents within a delimited area using highly polymorphic markers, such as nuclear microsatellites, to reconstruct dispersal events (Bode et al., 2018; Gerber et al., 2003). Parentage analyses directly estimate dispersal distance distributions, identify reproductive individuals, and provide estimates of their reproductive success. These direct methods rely on exhaustive or quasi-exhaustive sampling of reproductive individuals in a study plot and high-quality multilocus genotypes. For example, CERVUS, based on likelihood-based parentage assignment (Marshall et al., 2003), and COLONY, using a full-pedigree Bayesian framework (O. R. Jones & Wang, 2010), both permit the assignment of specific parents to offspring. When combined with maternally inherited markers such as plastid microsatellites (cpSSRs) or mitochondrial DNA (mtDNA)(maternally inherited in most angiosperms), maternal vs. paternal contributions can be distinguished (Ashley, 2010; Gerber et al., 2003; Wang, 2012). Spatially explicit mating models such as MEMM (Klein et al., 2008), NMπ2 (Chybicki, 2018) or MEMMseedlings (Oddou-Muratorio & Klein, 2008) allow to estimate a variety of mating system variables such as seed and pollen dispersal kernels, seed and pollen immigration rates into the study plot, and male and female fecundities and their drivers. These models require rich datasets, e.g. including individual and neighbourhood variables, that may be difficult to collect, e.g., in tropical ecosystems where tree species typically occur at low densities. The large stature of canopy tree species, their irregular and infrequent flowering, and the low density of reproductive individuals pose major obstacles to direct monitoring of gene flow (Bawa, 1990; Dick et al., 2008; Robledo-Arnuncio et al., 2014; Ward et al., 2005). The difficulty of sampling both adult trees and offspring across sufficiently large areas also complicates direct assessments of gene dispersal. Empirical studies in the Amazon rainforest remain scarce and often spatially limited, leading to significant gaps in our understanding of how gene flow operates in different tree species and across diverse tropical environments (Dick et al., 2008; Gerber et al., 2003; Monteiro et al., 2019). Topographic variation, forest disturbance, and differences in population density have been shown to influence SGS intensity across and within species (Sebbenn et al., 2008; Torroba-Balmori et al., 2017; Ward et al., 2005). Combining indirect (SGS-based) and direct, parentage-based approaches can offer complementary perspectives (Major et al., 2021; Oddou-Muratorio & Klein, 2008): SGS showcases the memory of past dispersal and demography, while parentage captures the fine-scale ecological processes of the current generation.

When estimates from direct methods and indirect methods converge, this typically suggests a relatively stable population dynamic over several generations, e.g., arising from demographic continuity and landscape stability (Dunphy et al., 2004; Gargiulo et al., 2023; Oddou-Muratorio & Klein, 2008; Vandergast et al., 2007). Conversely, discrepancies between direct and indirect estimates often signal recent ecological perturbations. Disturbances can rapidly disrupt gene flow without immediately altering the SGS, which integrates gene flow over longer time periods (Dutech et al., 2005; Gargiulo et al., 2024). The temporal lagging of dispersal estimates from SGS means that they may still reflect pre-disturbance patterns, while direct estimates reveal recent shifts in reproductive success, connectivity, or dispersal distances (Robledo-Arnuncio et al., 2014). Comparing gene flow estimates from these two approaches therefore enables the detection of hidden demographic changes: for example, logging may decrease the number of effective pollen or seed dispersers, affecting real-time connectivity, but adult trees’ SGS patterns might still reflect the more continuous gene flow of previous generations (Bacles et al., 2006; Oddou-Muratorio & Klein, 2008). In some cases, compensatory mechanisms such as increased reproductive success of remaining individuals or changes in disperser behaviour may maintain effective gene flow despite anthropogenic disturbance (Sebbenn et al., 2008). Comparing indirect and direct estimates of dispersal, along with estimates of reproductive success, is thus a more complete approach for identifying the degree and timing of demographic shifts. This approach allows testing hypotheses about disturbance intensity, resilience capacity, and the timescale of demographic recovery, offering valuable insights for the management and conservation of tree populations in increasingly human-impacted tropical landscapes.

Here, we aimed to better understand the dynamics of historical and contemporary gene flow in *Dicorynia guianensis* Amshoff (Fabaceae), a common tree species typical of tropical rainforests in French Guiana. We compared direct and indirect approaches to characterize seed and pollen dispersal in four forest sites representing natural populations that differ in ecological and anthropogenic context. Specifically, we aimed to: 1) Characterize patterns of genetic diversity and spatial genetic structure in four sites using highly polymorphic nuclear and plastid microsatellite markers; 2) Analyze SGS in two diameter size classes corresponding to functional categories (non-reproductive seedlings - saplings - subadults and potentially reproductive adults) to detect potential shifts related to recruitment and demographic filtering; 3) Compare direct contemporary and indirect historical dispersal metrics across ecological contexts; 4) Quantify and compare individual reproductive success across sites, to evaluate how reproductive output is distributed among individuals in different contexts; 5) Determine whether combining parentage analysis and SGS-based gene dispersal estimates with reproductive success data can serve as an early-warning framework to detect incipient losses of genetic diversity and adaptive potential in tropical tree populations (Bonnier et al., 2023; Gargiulo et al., 2024; Major et al., 2021). By linking ecological and evolutionary time scales, our approach underscores the value of combining diverse analytical methods to achieve the most accurate assessment of population demographic and genetic conservation status.

## 2. Material and method

### 2.1. Study species

*Dicorynia guianensis* (Fabaceae), known as angelique, is an Amazonian endemic tree species found commonly in *terra firma* forests of French Guiana and Suriname, and rarely in Guyana (Falcão et al., 2022). In French Guiana, *D. guianensis* is ecologically and economically important; it represents 54% of the timber production of the territory and its wood is used locally in carpentry, joinery, flooring, and harbour works (Flora, 2018; Guitet et al., 2014). In terms of wood productivity and biomass, it is considered the 16th hyperdominant species in the Amazon basin (Fauset et al., 2015). *D. guianensis* can reach heights of 45–50 meters and diameters exceeding 100 cm. The flowers are asymmetric with a reduced perianth, and notably, an upper and a lower stamen that are both polysporangiate. Pollination is likely insect-mediated, involving large bees, with flowers emitting a strong scent during early morning anthesis (Jésel, 2005). The species is predominantly outcrossing, with pollen dispersal reaching at least 190 m around the source tree (Caron et al., 1998). The fruits of *D. guianensis* are samaroid, indehiscent, and adapted for wind dispersal within generally limited dispersal distances (Falcão et al., 2022) typically up to 30 m, with only 5% of seeds reported to disperse beyond 50 m (Caron et al., 1998). Seeds are exposed to pre- and post-dispersal predation by insects, rodents, and parrots and occasional secondary seed dispersal by animals has been hypothesised (P. Forget, 1988; Jésel, 2005). The species is primarily distributed in genetically differentiated aggregates (Caron et al., 1998; Latouche-Hallé et al., 2003), typically covering less than one hectare, with mature tree densities ranging from 4 to 18 per hectare (Bariteau, 1993; Gourlet-Fleury et al., 2004; PRFB, 2019; authors’ personal observation). *D. guianensis* populations from western, coastal French Guiana are genetically differentiated from those in the east and more inland, and there is variable, weak spatial genetic structure within populations (Bonnier et al., 2023, 2025; Caron et al., 2000).

### 2.2. Sampling and study sites

*D. guianensis* tissue samples were collected during several weeks of intensive sampling between 2022 and 2023, conducted in four natural populations across French Guiana: Sparouine (05°15’45”N, 54°10’10”W), Paracou (05°15’30”N, 52°56’16”W), Nouragues (04°04’55”N, 52°40’12”W) and Regina (04°04’42”N, 52°05’23”W)(Fig. 1). The study sites are characterized by variable densities of *D. guianensis* (Table 1), contrasted climatic and pedologic conditions, and variable levels of anthropogenic disturbance, allowing us to study spatial genetic structure and dispersal patterns of *D. guianensis* in different environments (Guitet, Olivier, et al., 2015; Guitet, Pélissier, et al., 2015). Sparouine is located in western French Guiana, this area is sparsely populated and relatively difficult to access, which likely limits hunting pressure and other direct human disturbances. Nouragues is located within a strictly protected natural reserve in central French Guiana. The site is accessible only after several hours of river transport or by helicopter, and entry is regulated through official authorization. Human presence is therefore minimal, and the forest structure remains largely undisturbed in recent decades. Paracou is situated near the coastal region and is directly accessible via the RN1 road, in proximity to the majority of urban centers and human population in French Guiana (Voss et al., 2001). This accessibility facilitates human activities, including subsistence and recreational hunting. Local communities commonly hunt a wide range of vertebrates, from ungulates to small frugivorous birds. Regina underwent selective logging in 2012, approximately 12 years prior to sampling. Within the studied plot, 42 trees of *D. guianensis* with DBH ranging from 60 to 115 cm were harvested. Logging operations, including tree felling, skid trails, and log landing areas, resulted in approximately 2.1 hectares of canopy openings within the study area (12%). In addition to modifying local canopy structure and adult tree density, the creation of forest roads increased accessibility to previously isolated forest areas, potentially facilitating human entry and hunting activities. The sampled sites at Sparouine (30 ha), Paracou (25 ha, permanent plot number 16, Derroire et al., 2025), Nouragues (38 ha), and Regina (17 ha) were systematically surveyed for *D. guianensis* individuals ≥1 cm Diameter at Breast Height (DBH), aiming for an as complete as possible sampling of individuals. The GPS position, DBH, and sampling altitude of each tree were recorded during sampling. Leaf or cambium samples were collected and stored in silica gel at air-conditioned room temperature until DNA extraction. A total of 1,528 individual samples were collected: Sparouine (367), Paracou (360), Nouragues (383), Regina (417) (Fig. 1). For analysis requirements, the trees studied were divided into two size classes defined according to their DBH. Trees with DBH under 30 cm were classified as seedlings-saplings-subadults (SSS), and those with DBH over 30 cm as adults (ADL). Seedlings-saplings-subadults trees correspond to individuals that do not have the ability to contribute to the reproductive event in the studied plots, while trees with DBH ≥ 30 cm regularly bear fruit and are thus considered as potentially reproductive adults (*field observation*).

**Table 1.**
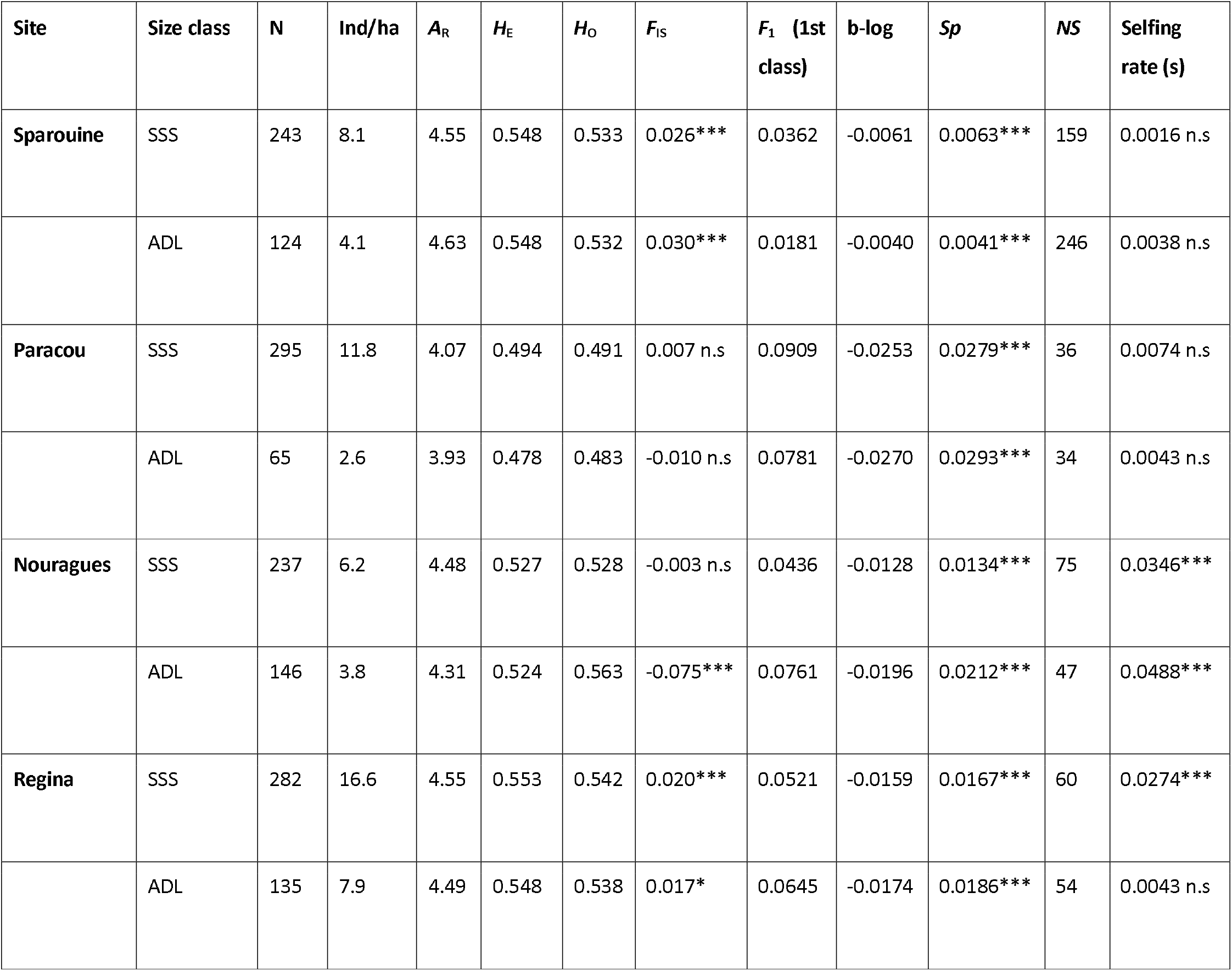
Genetic diversity and spatial genetic structure parameters for *Dicorynia guianensis* individuals categorized into two size classes: Seedlings-saplings-subadults (SSS), DBH < 30 cm and Adults (ADL), DBH > 30 cm across four study sites in French Guiana. N, number of individuals sampled; *A*_R_, allelic richness standardized to 50 gene copies; *H*_E_, expected heterozygosity; *H*_O_, observed heterozygosity; *F*_IS_, inbreeding coefficient, and significance of permutation test (***, P < 0.001; *, P < 0.05; ns, not significant); *F*_i_ (1st class), average pairwise kinship coefficient within the first distance class; b-log, regression slope of *F*_i_ on the logarithm of distance; *Sp*, strength of spatial genetic structure and significance (***, P < 0.001); *NS*, Wright’s neighbourhood size; selfing rate (s), and jackknife significance (***, P < 0.001; ns, not significant).

| Site | Size class | N | Ind/ha | $A_R$ | $H_E$ | $H_O$ | $F_{IS}$ | $F_1$ (1st class) | b-log | $Sp$ | NS | Selfing rate (s) |
| --- | --- | --- | --- | --- | --- | --- | --- | --- | --- | --- | --- | --- |
| Sparouine | SSS | 243 | 8.1 | 4.55 | 0.548 | 0.533 | 0.026*** | 0.0362 | -0.0061 | 0.0063*** | 159 | 0.0016 n.s |
|  | ADL | 124 | 4.1 | 4.63 | 0.548 | 0.532 | 0.030*** | 0.0181 | -0.0040 | 0.0041*** | 246 | 0.0038 n.s |
| Paracou | SSS | 295 | 11.8 | 4.07 | 0.494 | 0.491 | 0.007 n.s | 0.0909 | -0.0253 | 0.0279*** | 36 | 0.0074 n.s |
|  | ADL | 65 | 2.6 | 3.93 | 0.478 | 0.483 | -0.010 n.s | 0.0781 | -0.0270 | 0.0293*** | 34 | 0.0043 n.s |
| Nouragues | SSS | 237 | 6.2 | 4.48 | 0.527 | 0.528 | -0.003 n.s | 0.0436 | -0.0128 | 0.0134*** | 75 | 0.0346*** |
|  | ADL | 146 | 3.8 | 4.31 | 0.524 | 0.563 | -0.075*** | 0.0761 | -0.0196 | 0.0212*** | 47 | 0.0488*** |
| Regina | SSS | 282 | 16.6 | 4.55 | 0.553 | 0.542 | 0.020*** | 0.0521 | -0.0159 | 0.0167*** | 60 | 0.0274*** |
|  | ADL | 135 | 7.9 | 4.49 | 0.548 | 0.538 | 0.017* | 0.0645 | -0.0174 | 0.0186*** | 54 | 0.0043 n.s |

**Figure 1.**
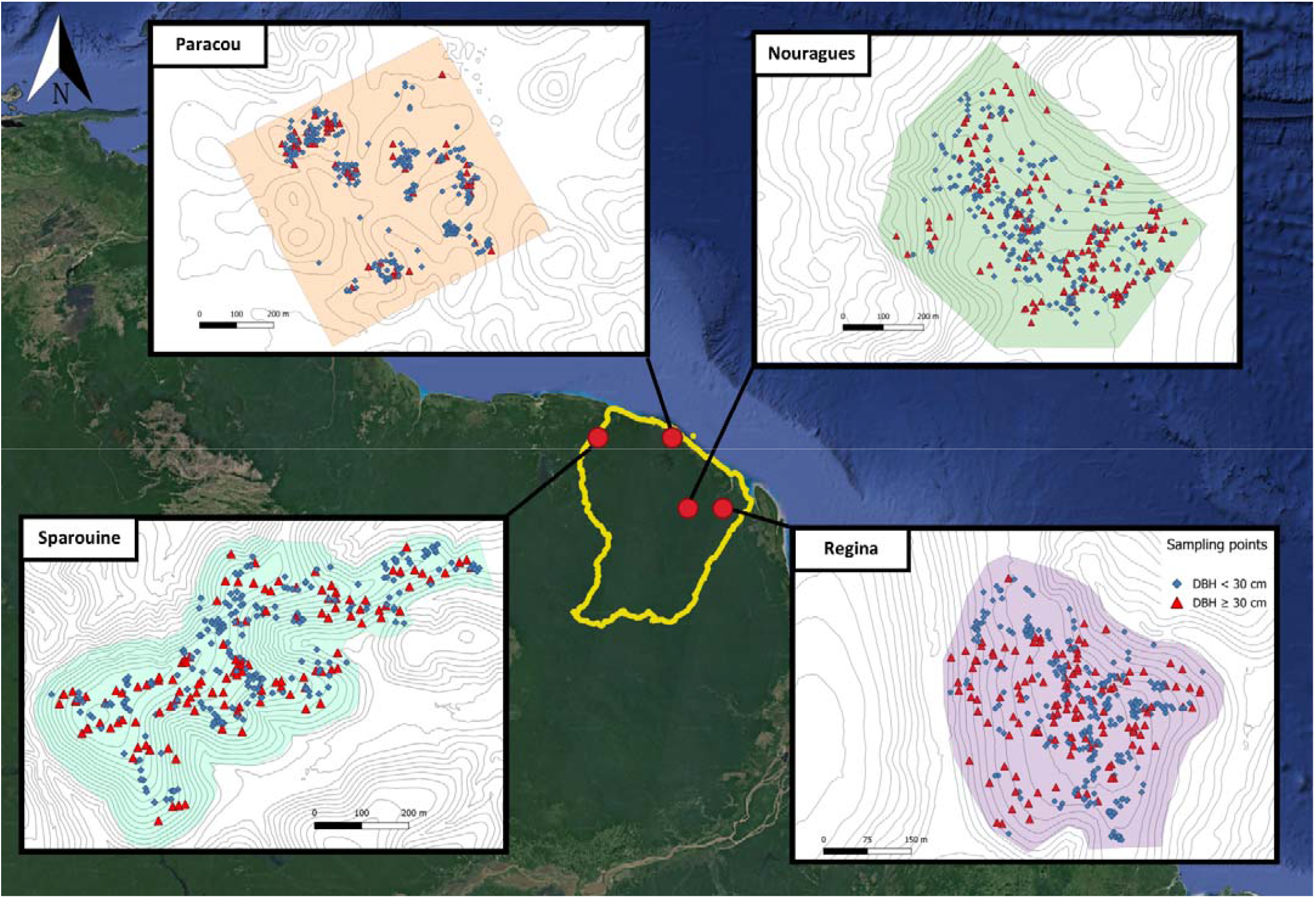
Geographic location and spatial distribution of *Dicorynia guianensis* individuals across four study sites in French Guiana. Map showing the position of the four sampled forest sites (Sparouine, Paracou, Nouragues, and Regina) within French Guiana (yellow boundary) in Northern South America. Sampling points are colour-coded by diameter at breast height (DBH): seedlings-saplings-subadults (DBH < 30 cm, blue rhombuses) and adults (DBH ≥ 30 cm, red triangles). Coloured shaded areas represent the sampling zone for each plot. Grey contour lines represent topographic isolines at 5 m intervals.

### 2.3. Regional genetic structure and genetic diversity

The genetic structure and diversity of *Dicorynia guianensis* populations was assessed at 66 nuclear microsatellites markers (SSRs) and 23 plastid loci (SSRs or single nucleotide polymorphisms, SNPs) at the four study sites. DNA extraction followed a CTAB protocol and microsatellite marker design, genotyping-by-sequencing and marker quality control followed the method described in (Lepais et al., 2020)(details in *Supporting information 1*). Genetic clustering was inferred using STRUCTURE software (Evanno et al., 2005) to assess the consistency of genetic cluster membership of individuals across the four sites. The analysis used an admixture model with correlated allele frequencies and assessed a number of genetic clusters (K) from 1 to 10, using a burn-in period of 50,000 iterations followed by 500,000 Markov Chain Monte Carlo (MCMC) repetitions for each run. The results were analyzed using STRUCTURE Harvester (Earl & vonHoldt, 2012) to determine the most likely number of clusters and to evaluate the individual probabilities of assignment to the inferred clusters.

The inter-sites and intra-sites genetic diversity of *D. guianensis* was assessed across the four study sites using SPAGeDi software (Hardy & Vekemans, 2002). Genetic diversity indices were calculated for ADL and SSS for each site. This analysis included the computation of pairwise *F*_*ST*_ values to quantify genetic differentiation among sites. Genetic diversity indices were calculated, including allelic richness standardized to 50 gene copies (*A*_*R*_), observed heterozygosity (*H*_*O*_), expected heterozygosity (*H*_*E*_), and inbreeding coefficient (*F*_*IS*_). Significance was assessed by permutation tests performed with 10,000 randomisations of gene copies among individuals. Selfing rate (s) was also computed with SPAGeDi (Hardy & Vekemans, 2002). The significance of selfing rate estimates was assessed using the multilocus identity disequilibrium parameter g_2_, with P-values obtained via jackknife over loci (Hardy et al., 2006). For genetic diversity at plastid markers, allelic richness (*A*_*R*_), expected heterozygosity (*H*_*E*_), and the effective number of alleles were also computed.

### 2.4. Spatial genetic structure within plots and historical dispersal parameters

Spatial principal component analysis (sPCA) was done using the spca() function in the adegenet R package (Jombart, 2008) to detect, test, and visualize spatial patterns of genetic variability within plots. The global test (G-test) was used to assess the significance of overall genetic structure within plots and the local test (L-test) to identify significant genetic repulsion between neighbours. Spatial interpolation of lagged sPCA scores was displayed in maps to illustrate and facilitate the interpretation of the spatial distribution of genetic variability. To test the link between genetic structure at site level and topography, a simple linear regression model was used to estimate the effect of altitude on lagged scores using the lm() function in R.

To study the decay of genetic similarity with spatial distance, spatial genetic structure (SGS) was assessed by plotting pairwise relatedness coefficients (*F*_*ij*_, Loiselle et al., 1995) against the logarithm of pairwise geographic distance using six predefined upper limits of the distance classes: 30, 60, 90, 130, 170, 220, 300, and 600 m. This definition of distance classes considered the previously estimated dispersal distances of *D. guianensis* seeds and pollen (Caron et al., 1998; P. Forget, 1988; Jésel, 2005), while aiming for similar sample sizes in each class. The average *F*_*ij*_ of the first distance class, *F*_*1*_, was used as an estimate of within-aggregate relatedness. The strength of SGS was quantified using the *Sp* statistic, defined as *Sp=-b*_*log*_*/(1-F*_*1*_), where *b*_*log*_ is the regression slope of *F*_*ij*_ on the logarithm of distance (Vekemans & Hardy, 2004). These parameters were computed for both nuclear and plastid markers. As a proxy of local effective population size, Wright’s neighbourhood size (*NS*) was calculated as *NS=1/Sp* (Wright, 1943) on nuclear SSRs, providing an estimate of the number of individuals effectively contributing to reproduction within a local area (Nunney, 2016). For plastid markers, the same framework was used to estimate seed dispersal distance.

Under the assumption of drift-dispersal equilibrium, *Sp* is expected to approximate 1/(4πD⍰ σ_g_^2^) for nuclear markers and 1/(2πD⍰σ_s_^2^) for plastid markers, where σ_g_^2^ and σ_s_^2^ represent half the mean squared dispersal distances of genes and seeds (Born et al., 2008; Vekemans & Hardy, 2004). Using this model, we estimated historical gene and seed dispersal distances (σ_g_, σ_s_) for each site. Effective population density (De) was estimated as 25% of the census density of adult individuals (DBH ≥ 30 cm) in each site, considering that ADL have variable age-related mortality and fertility rates. This reflects field observations of *D. guianensis*, which show marked interannual variation in flowering and seed production, limiting the number of adults contributing to reproduction each year. The estimation of σ_g_ and σ_s_ was conducted via an iterative procedure of the kinship-ln(distance) regression, in order to estimate these statistics from the appropriate distance range (Heuertz et al., 2003). Pollen dispersal distances (σ_p_) were inferred using the equation σ_g_^2^ = σ_s_^2^ + ½σ_p_^2^ (Crawford, 1984), allowing the computation of the pollen-to-seed dispersal ratio (σ_p_/σ_s_) and the seed-to-genes dispersal ratio (σ_s_/σ_g_).

### 2.5. Parentage analysis

Parentage was analysed using CERVUS v3.0.7 software (Marshall et al., 2003). CERVUS uses a likelihood-based approach to identify candidate parents for each potential offspring (seedlings - saplings - subadults). We conducted simulations (n=10,000 offspring) to establish critical thresholds for parentage assignment confidence based on allele frequencies. Simulation parameters included 500 candidate parents, a parent sampling rate of 25% and allowing polygamy in both sexes. Only parent-offspring assignments reaching relaxed (80%) confidence thresholds based on log-likelihood ratio were retained for subsequent analyses. Each analysis was replicated 10 times to ensure consistency of results. Parentage analysis with CERVUS provided a candidate pair of parents for subsequent analyses. We also used the COLONY software (O. R. Jones & Wang, 2010) in full likelihood mode, the most accurate mode as verified by simulated and empirical data analyses (Wang, 2012). Key assumptions included monoecious diploid individuals, and potential polygamy in both sexes. The size of the candidate parental population was set to 25%, similar to analyses in CERVUS. COLONY outputs (BestConfig and BestCluster files) provided parent-offspring assignments and allowed identification of full- and half-sibling families, facilitating further family structure analysis. Identified family clusters were plotted on site maps using the R package ggplot2 (Wickham, 2009), presenting only clusters composed of more than 5 individuals for clarity. Results from CERVUS and COLONY were merged, considering as robust only matching assignments between the two methods. For each offspring with an assigned parent pair, maternal identity was inferred based on plastid haplotype sharing between parent and offspring, provided a distinct haplotype was observed for the second parent, inferred as the biological father. Offspring with multiple matching mothers were excluded. Distances between parents and offspring, mean and median seed dispersal for mothers and pollen dispersal for fathers were calculated for each site. To assess uncertainty around mean estimates, 95% confidence intervals were calculated using non-parametric bootstrap resampling with 10,000 iterations. Data manipulation and cleaning steps used dplyr (Wickham et al., 2023) and tidyr (Wickham et al., 2014), and geographic distance calculations utilized the geosphere package (Hijmans, 2010).

### 2.6. Reproductive success

Based on matching parentage assignments between CERVUS and COLONY, we conducted a series of complementary analyses on reproductive success. We assessed the relationship between reproductive success and tree size by site performing linear regressions using the lm() function in R, testing the effect of DBH on the number of offspring assigned per tree. We also examined the distribution of reproductive roles (mother vs. father) for each parent. A paired Wilcoxon signed-rank test (Woolson, 2005) was used to compare the number of offspring produced as mothers versus fathers and DBH values of parents were compared by reproductive roles. To quantify reproductive inequality within each site, we computed the Gini index using the *Gini()* function from the R package *ineq* (Zeileis & Kleiber, 2014), based on the total number of offspring assigned to each parent. The Gini index, a continuous measure of inequality ranging from 0 to 1 offering insight into reproductive skew within each site (Gastwirth, 1972). Values above 0.45 are typically interpreted as reflecting substantial skew in reproductive success (Ross et al., 2020, 2023). Significant differences in reproductive skew across sites were tested using 10,000 permutations of individual reproductive success values across sites by randomly reassigning individual reproductive outputs across sites. Parent-offspring clusters inferred by COLONY were used to characterize family group structure within sites and to assess the variance in reproductive success among clusters (O. R. Jones & Wang, 2010).

## 3. Results

### 3.1. Genetic structure and diversity

All sampled sites were genetically differentiated and formed distinct genetic clusters, with no evidence of recent migrants among sites (Supporting Information 2). Allelic richness (A_R_) was lowest in Paracou (*A*_*R*_ = 3.93, 4.07) and highest in Regina (*A*_*R*_ = 4.49, 4.55) and Sparouine (*A*_*R*_ = 4.55, 4.63), (Table 1). When comparing size classes (<30 cm DBH vs ≥30 cm DBH), allelic richness and expected heterozygosity remained broadly similar within each site, with no evidence of marked erosion of genetic diversity between size classes. Seedlings-saplings-subadults (SSS) individuals tended to show slightly higher *A*_*R*_ than adults in Paracou and Nouragues, whereas differences were minimal in Sparouine and Regina.

Expected heterozygosity (*H*_*E*_) followed a similar pattern, ranging from 0.478 in Paracou adults to 0.553 in Regina SSS individuals. Inbreeding coefficients (*F*_*IS*_) were generally low but site-dependent. Significant positive *F*_*IS*_ was found in Sparouine in SSS (*F*_*IS*_ = 0.026***) and in ADL (*F*_*IS*_ = 0.030***) individuals, and in Regina in SSS (*F*_*IS*_ = 0.020***) and in ADL (*F*_*IS*_ = 0.017*) individuals. Paracou had non-significant *F*_*IS*_ while Nouragues displayed a strong heterozygote excess in adults (*F*_*IS*_ = −0.075***). Selfing rates were generally low across sites with an average value of 0.009.

Plastid markers resolved 27 haplotypes across the four sites, with 5 to 13 haplotypes per site slightly lower plastid genetic diversity in Paracou (*H*_*E*_ = 0.512) than in other sites (0.566 in Sparouine, 0.555 in Regina, and 0.548 in Nouragues) (*Supporting information 2*, Table S2.3).

### 3.2. Spatial genetic structure and historical dispersal estimates

The sPCA analysis revealed significant global genetic structures (P ≤ 1e-06) in all sites. Spatial genetic structure gradients illustrated in lagged score maps appeared to be visually correlated with altitude *(Supporting information 3*, Figure S3.1). This observation was supported by significant linear relationships between lagged scores and altitude in all sites (*Supporting information 3*, Figure S3.2).

The Sp statistic revealed variation in spatial genetic structure between size classes and among sites (Table 1; Figure 2). In Paracou, high *Sp* values were observed in both size classes, with *Sp* = 0.0279*** in SSS individuals and Sp = 0.0293*** in adults, associated with low neighbourhood sizes (NS = 36 and 34, respectively). *Sp* values were higher in SSS than in ADL individuals in Sparouine only. In Sparouine, neighbourhood size (*NS*) was higher in ADL (*NS* = 246) than in SSS individuals (*NS* = 153), consistent with a weaker *Sp* in the ADL class; a similar pattern was observed in Nouragues reflecting the sharp reduction in SGS intensity in mature trees.

**Figure 2.**
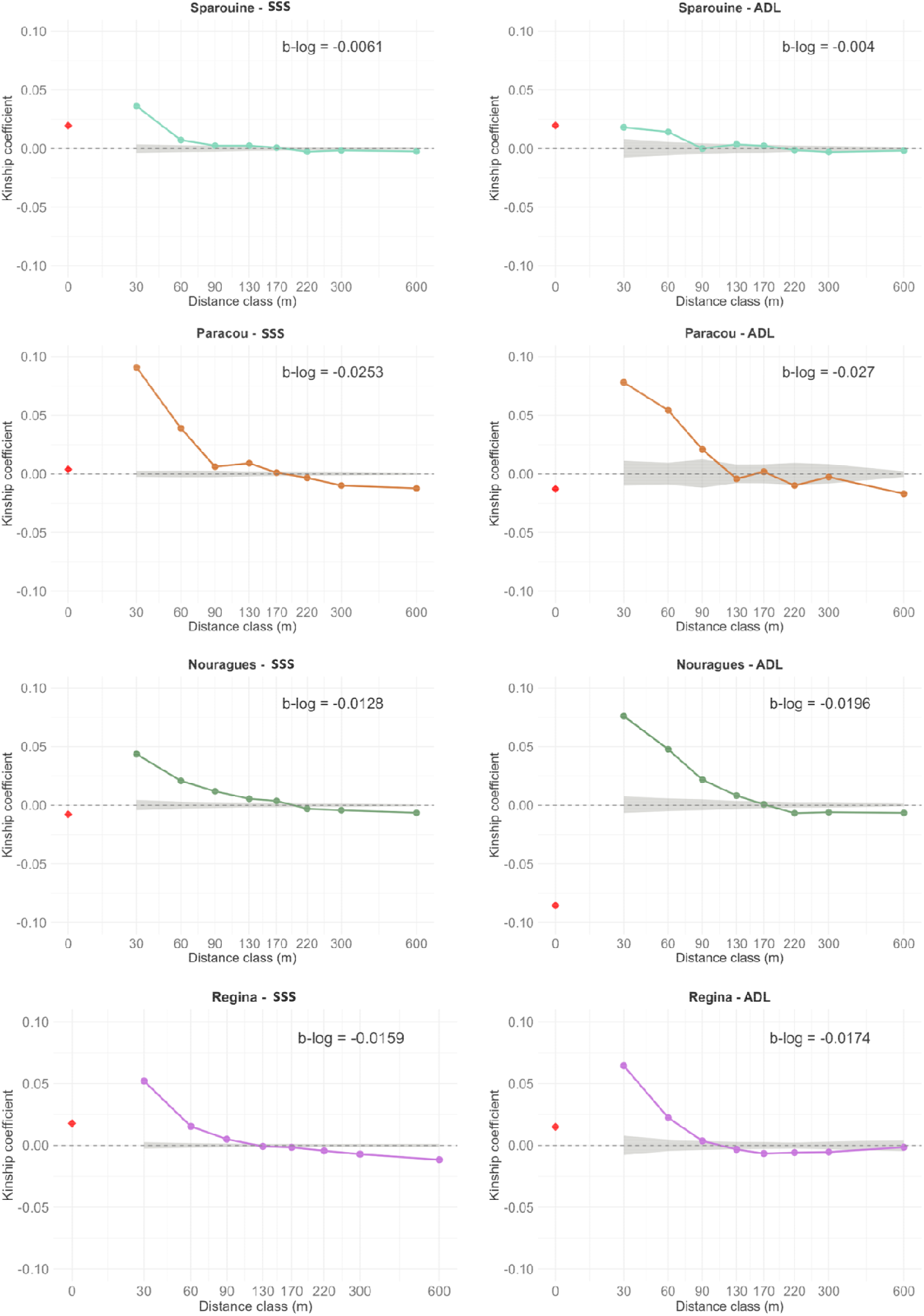
Spatial autocorrelograms of the kinship coefficient (*F*_ij_) as a function of geographic distance for the four *D. guianensis* study sites for adult and Seedlings-saplings-subadults size classes. The dashed grey area represents the 95% confidence interval for spatial randomness of genotypes, generated by 10,000 permutations of individual locations. The regression slope of kinship on log(distance) (b-log) is indicated for each site. The red rhombus represents the intra-individual kinship coefficient.

When spatial genetic structure was estimated using plastid markers, *Sp* values were consistently higher than those estimated from nuclear markers in all sites (Table 2). The highest plastid *Sp* value was observed in Regina (Sp = 0.285***). Historical gene dispersal distances (σ_g_) inferred from nuclear SGS varied among sites and were highest in Sparouine (σ_g_ = 433.4 m), followed by Paracou (σ_g_ = 204.4 m), Nouragues (σ_g_ = 197.7 m) and Regina (σ_g_ = 146.8 m). Decomposition into maternal and paternal components indicated that pollen dispersal distances (σ_p_) were clearly higher in Sparouine (σ_p_ = 574.4 m) compared to the other plots. Conversely, seed dispersal was much lower in Regina (σ_s_ = 53.12 m) than in the other sites (Table 2).

**Table 2.** Nuclear and plastid spatial genetic structure metrics and estimated historical dispersal distances in four *D. guianensis* study sites. *F*_1_, average pairwise kinship coefficient within the first distance class for nuclear (nucl) or plastid (cp) markers; b-log, regression slope of *F*_ij_ on the logarithm of distance for nuclear (nucl) or plastid (cp) markers; *Sp*, strength of spatial genetic structure (ADL) for nuclear (nucl) or plastid (cp) markers and significance (*** P < 0.001); σ_g_, σ_s_, and σ_p_ correspond to historical gene, seed, and pollen dispersal distances (in meters), respectively, estimated from SGS parameters.

| Sites | $F_1$ (nucl) | $b_{log}$ (nucl) | $Sp$ (nucl) | $F_1$ (cp) | $b_{log}$ (cp) | $Sp$ (cp) | $\sigma_g$<br>(m) | $\sigma_s$<br>(m) | $\sigma_p$<br>(m) |
| --- | --- | --- | --- | --- | --- | --- | --- | --- | --- |
| Sparouine | 0.025 | -0.004 | 0.004*** | 0.362 | -0.043 | 0.067*** | 433.4 | 151.1 | 574.4 |
| Paracou | 0.100 | -0.024 | 0.029*** | 0.440 | -0.076 | 0.136*** | 204.4 | 134.2 | 218.0 |
| Nouragues | 0.056 | -0.015 | 0.021*** | 0.495 | -0.075 | 0.149*** | 197.7 | 105.4 | 236.5 |
| Regina | 0.052 | -0.016 | 0.019*** | 0.561 | -0.125 | 0.285*** | 146.8 | 53.1 | 193.6 |

### 3.3. Parentage and contemporary gene flow

Parentage analyses combining CERVUS and COLONY, refined by chloroplast haplotype information, enabled robust reconstruction of parent-offspring relationships. Assignment success varied both by method and by site, reflecting differences in sampling completeness and local reproductive dynamics (Table 3, Fig. 3).

**Table 3.** Summary of parentage assignment results by software and maternal identification per site. Density (ADL), density of individuals with dbh ≥ 30cm per hectare; Number of potential parents (ADL) and Number of potential offsprings (SSS), sample sizes of the corresponding size classes; the proportion of offspring assigned by each software, the proportion of concordant assignments between both methods, the percentage of offspring with a genetically confirmed mother, based on cpSSR haplotypes and mean seed and pollen dispersal distances in meters ± half-width of the 95% confidence interval, calculated via non-parametric bootstrap (10,000 resamples).

| Site | Density<br>(ADL) | Number of<br>potential<br>parent<br>(ADL) | Number of<br>potential<br>offspring<br>(SSS) | Offspring<br>inferred by<br>CERVUS | Offspring<br>inferred by<br>COLONY | Common<br>offspring<br>inferred | Offspring<br>with<br>mother<br>identified | Mean<br>seed<br>dispersal<br>distance<br>(m) | Mean<br>pollen<br>dispersal<br>distance<br>(m) | Median<br>seed<br>dispersal<br>distance<br>(m) | Median<br>pollen<br>dispersal<br>distance<br>(m) |
| --- | --- | --- | --- | --- | --- | --- | --- | --- | --- | --- | --- |
| Sparouine | 4.1 | 124 | 202 | 14.4% | 27.1% | 13.7% | 6.9% | 100(±<br>60) | 293(± 65) | 40 | 283 |
| Paracou | 2.6 | 65 | 191 | 91.3% | 59.6% | 49.5% | 22.0% | 39 (± 21) | 220(± 37) | 21 | 188 |
| Nouragues | 3.8 | 146 | 295 | 59.4% | 47.6% | 42.8% | 21.4% | 91(± 18) | 203(± 29) | 74 | 176 |
| Regina | 7.9 | 135 | 202 | 90.2% | 77.3% | 75.5% | 38.6% | 41(± 12) | 135(± 15) | 21 | 132 |
| Overall | 13.8 | 470 | 890 | 60.8% | 51.3% | 43.6% | 21.4% | 62(± 11) | 205(± 16) | 34 | 188 |

**Figure 3.**
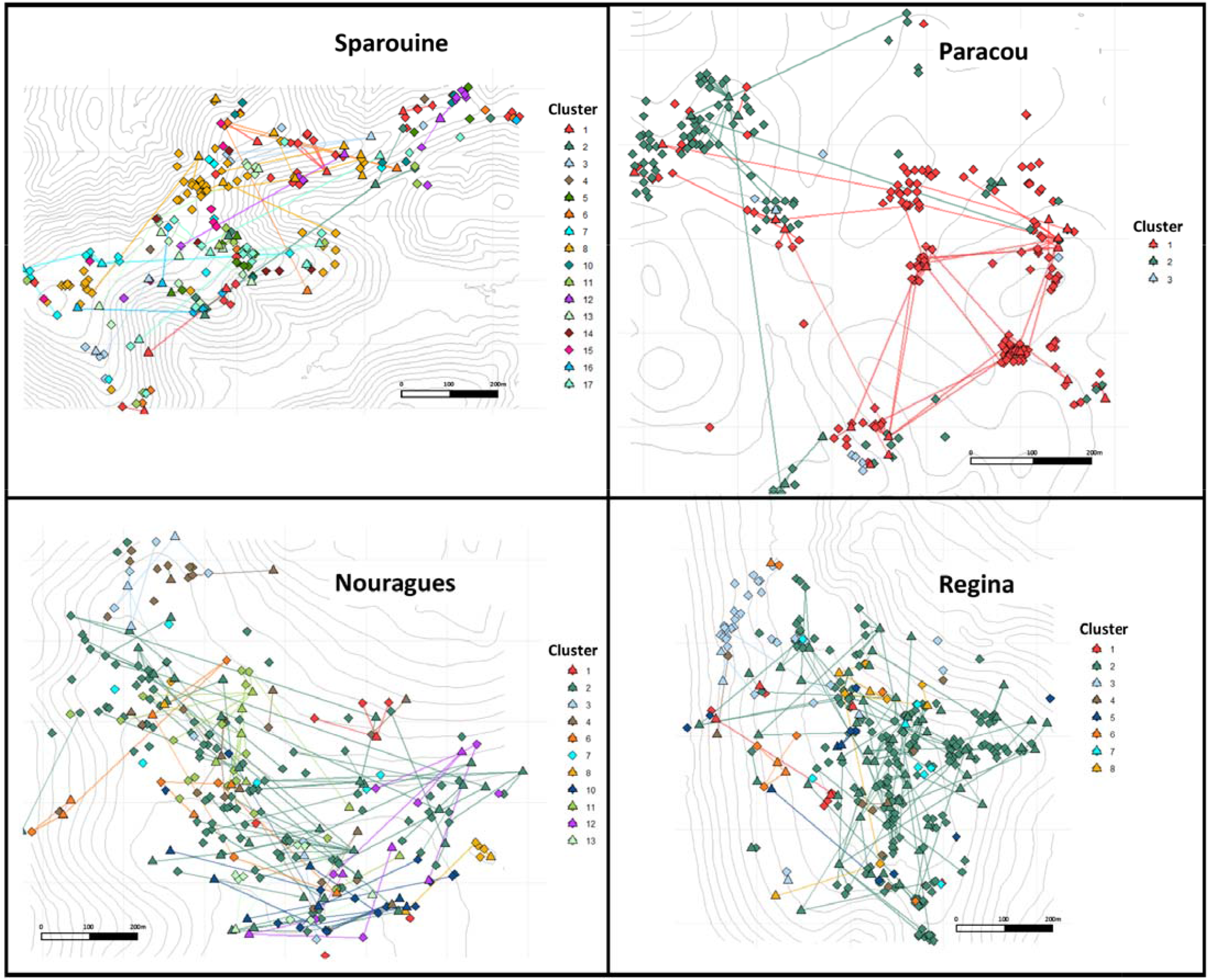
Spatial representation of reproductive clusters and parent-offspring relationships inferred using COLONY software in four *Dicorynia guianensis* populations. Only family clusters containing more than five individuals are shown. Each symbol represents a tree individual, colour-coded according to its assigned full-sibling family. Rhombuses represent offspring and triangles adults. Lines connect inferred parent-offspring pairs within each cluster.

The highest assignment rates were observed at Paracou (91.3% with CERVUS, 59.6% with COLONY) and Regina (90.2% with CERVUS, 77.3% with COLONY), while Sparouine showed the lowest values with only 14.4% of offspring assigned by CERVUS and 27.1% by COLONY. Concordance between the two methods also varied among sites: 43.6% of total offspring were assigned a parent pair by both tools. Incorporating plastid haplotype data further enabled the identification of mothers for 21.4% of these offspring, with the highest maternal assignment rate in Regina (38.6%), followed by Paracou (22.0%), Nouragues (21.4%), and Sparouine (6.9%). Mean and median seed and pollen dispersal distances inferred from mother and father assignments varied among sites (*Supporting information 4*, Figure S4.1). Median seed dispersal was shortest in Paracou and Regina (21 m), while much longer in Nouragues (74 m) and Sparouine (40 m) (*Supporting information 4*, Figure S4.2). In all cases, mean and median pollen dispersal distances were ca. 3-6 times greater than seed dispersal distances, with Sparouine again standing out with the longest mean pollen dispersal distance (293 m) and Regina displaying the shortest (135 m) (*Supporting information 4*, Figure S4.3). Categorizing individuals by family provided by COLONY enabled us to observe variations in the diversity of links among individuals present in the sites, with a total of 23, 5, 15, and 12 families respectively in Sparouine, Paracou, Nouragues, and Regina. Families of more than 5 individuals are shown on a map (Fig. 3).

Barcharts of seed and pollen dispersal distances across all four sites combined provide a detailed view of the overall dispersal dynamics in *Dicorynia guianensis* populations (*Supporting information 4*, Figure S4.4). Most seeds were dispersed within 50 meters of the maternal tree, with a sharp decline in frequency beyond this range. A few rare long-distance events were recorded, reaching up to nearly 400 meters. The mean and median seed dispersal distances (∼62 m and 34 m) reflect a short overall seed dispersal distance, with the mean distance driven by rare long-distance dispersal (Supporting information 4, Figure S4.4A). Pollen dispersal shows a peak around 150–200 meters and a median of 188 meters. In contrast to seed dispersal, pollen dispersal events are more evenly distributed across distances, and long-distance events (300–500+ meters) are relatively frequent (Supporting information 4, Figure S4.4B).

### 3.4 Variation in reproductive success

Linear regression analyses revealed a significant positive relationship between DBH and reproductive output (i.e., number of offspring assigned) in Paracou, Nouragues, and Regina (*Supporting information 4*, Figure S4.5). The paired Wilcoxon signed-rank tests revealed no significant difference in the number of offspring assigned to mothers versus to fathers in any site, indicating a balanced contribution of reproductive roles. When comparing the DBH of assigned mothers and fathers, only Nouragues showed a statistically significant difference, with mothers presenting larger DBH values than fathers (P ≤ 0.002). Values from the Gini index to quantify reproductive skew showed the highest levels of reproductive inequality in Paracou (Gini = 0.504) and Regina (Gini = 0.474), followed by Nouragues (Gini = 0.372). Sparouine exhibited a lower Gini index (0.206). Pairwise permutation tests confirmed that the difference in reproductive skew between Sparouine and each of the other sites was highly significant (p < 0.001), while pairwise comparisons between Regina, Paracou, and Nouragues were not significant.

Analysis of family clusters from COLONY highlighted contrasts in reproductive structure across sites (Figure 4, *Supporting information 4*, Figure S4.6). In Regina, cluster 2 included 193 offspring, and in Paracou, clusters 1 and 2 combined reached 270 offspring. Sparouine showed a much more balanced distribution, with 23 clusters and a maximum of 53 offspring per cluster, suggesting less reproductive dominance. Nouragues exhibited an intermediate pattern, with one large cluster of 117 offspring, but also several smaller ones.

**Figure 4.**
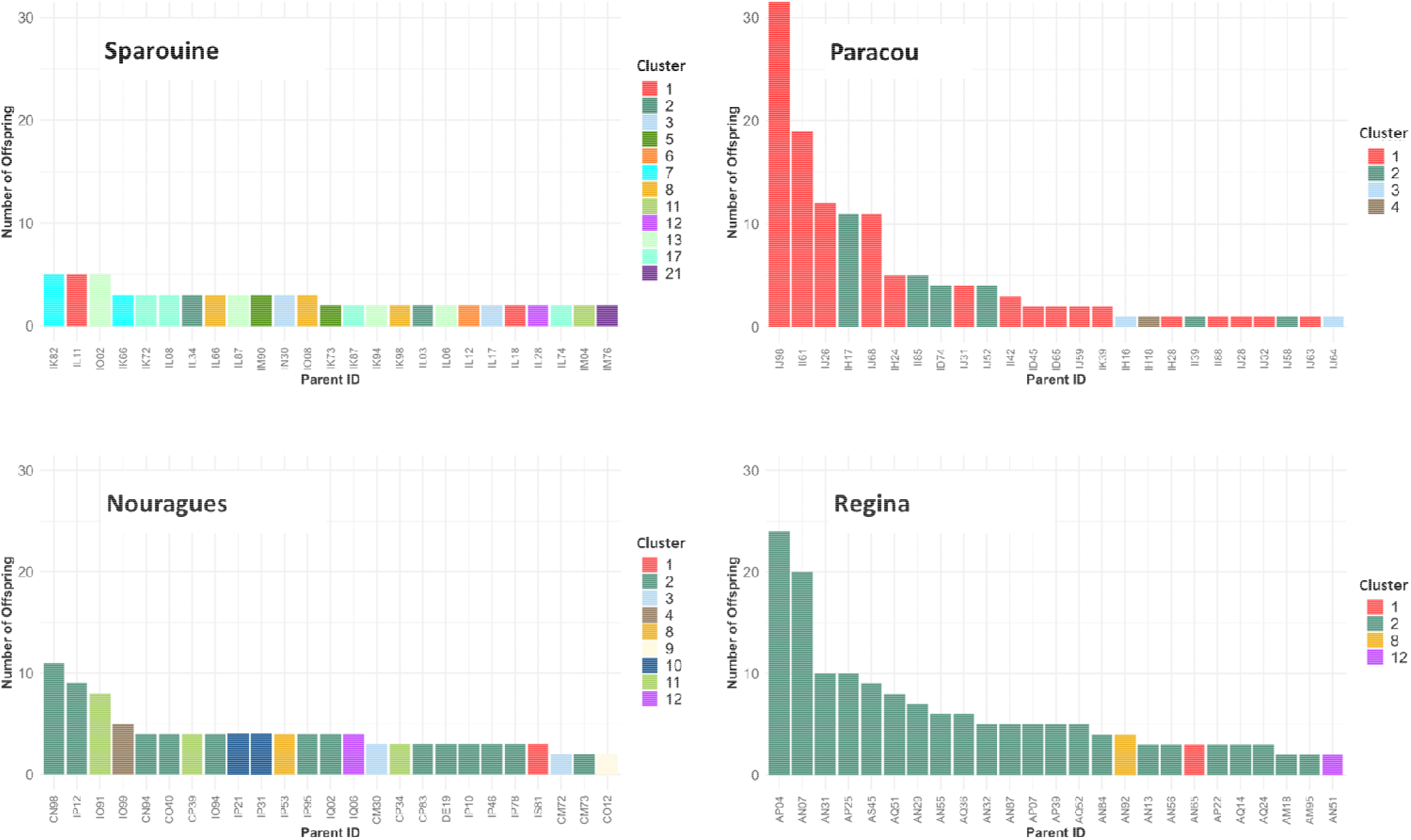
Individual reproductive success across four *Dicorynia guianensis* populations associated by cluster. Barplots showing the number of offspring assigned to the 25 most reproductively successful adult trees in each site (Sparouine, Paracou, Nouragues, and Regina), based on parentage assignments from COLONY. Each bar represents a parent individual (Parent ID), and bar colours indicate the reproductive cluster to which the individual belongs.

## 4. Discussion

### 4.1. Patterns of structure, genetic diversity and inbreeding

Spatial PCA confirmed significant global structure within all sites, while local structure remained limited. This indicates that genetic variation in sites is spatially structured on a large scale, meaning that individuals located closer to each other tend to be more genetically similar. Patterns of genetic variation within sites appeared to be shaped in part by elevation, with significant spatial correlations detected in three sites, a common phenomenon in species of neotropical trees linked to adaptation to micro-environmental conditions (Schmitt et al., 2021; Torroba-Balmori et al., 2017). These results highlight the importance of both local ecological factors and regional historical processes in shaping the genetic structure of Neotropical tree populations.

Patterns of genetic diversity revealed a spatial heterogeneity across the studied populations. Sparouine appeared to host the most genetically diverse population, while Paracou consistently displayed lower diversity across both nuclear and plastid markers. These contrasts suggest differences in historical demography, levels of connectivity and, possibly past disturbance regimes (Petit & Hampe, 2006; Stier et al., 2020). Despite this variation, heterozygosity levels remained relatively stable among populations, and inbreeding coefficients were low overall. This shows that for *D. guianensis*, despite sites with different contexts and histories, we do not observe any strong deviations from Hardy–Weinberg expectations. Finally, Nouragues had the highest selfing rate in adults (s = 0.0406) among the four sites, despite being located in a protected reserve expected to preserve natural dispersal dynamics and high outcrossing rates. Low anthropogenic disturbance alone does not guarantee reduced self-fertilization. In legumes tree species, mating systems are variable: many engage in mixed mating systems, allowing both outcrossing and selfing (Goncalves et al., 2019), while others show mechanisms of self-incompatibility (Delaney & Igić, 2022).

### 4.2. Size class variations: contrasting genetic structure across size classes

Spatial genetic structure differed between size classes across sites, but contrary to classical expectations, SGS was not systematically stronger in SSS individuals. With the exception of Sparouine, *Sp* values were equal or higher in adults than in SSS individuals (Table 1; Figure 2). This pattern contrasts with the commonly reported life-stage dynamic in tropical trees, where stronger SGS in juveniles reflects short-distance seed dispersal and localized recruitment near maternal trees, followed by progressive erosion of relatedness through density-dependent mortality and demographic thinning (Epperson & Alvarez-Buylla, 1997; Hardesty et al., 2005; Hardy et al., 2006; Sola et al., 2022). Although differences between size classes remain moderate in magnitude, they are a corollary of the strongly aggregative spatial dynamics of *Dicorynia guianensis* populations in French Guiana, already well documented at the ecological level (Guitet et al., 2014; Kokou, 1994). The persistence of strong SGS in adult size classes may reflect long-term spatial stability of recruitment rather than erosion of genetic clustering through demographic thinning. In species with limited seed dispersal and spatially aggregated regeneration niches, repeated recruitment in favourable microsites can maintain fine-scale genetic clustering across generations (Hardy et al., 2006; F. A. Jones & Hubbell, 2006). Moreover, if effective pollen dispersal only partially counteracts restricted seed dispersal, related individuals may remain spatially structured even at mature stages (Vekemans & Hardy, 2004).

Sparouine remains distinct, with weaker overall SGS compared to the other sites. This reduced spatial structuring is consistent with its larger neighbourhood size and higher inferred historical gene dispersal distances (Table 2). Together, these results suggest that size class-level SGS variation primarily reflects differences in site-specific demographic and spatial aggregation patterns rather than a uniform ontogenetic thinning process. Genetic diversity indices remained stable between size classes within each site (Table 1). Such patterns indicate that size classes-level changes in SGS are not accompanied by shifts in genetic diversity, and therefore likely reflect spatial aggregation dynamics rather than strong genetic erosion or size classes-specific bottlenecks (Ng et al., 2004; Seidler & Plotkin, 2006).

### 4.3. Spatial genetic structure as a legacy of demographic and dispersal processes

SGS integrates the cumulative effects of gene dispersal and demographic processes over multiple generations (Vekemans & Hardy, 2004), dispersal distances estimated from σ parameters reflect historical rather than strictly contemporary patterns of gene movement. Among all sites, Sparouine exhibited the highest historical gene dispersal distances, driven largely by high pollen dispersal distance while seed dispersal distances were closer to those observed in the other plots. This pattern highlights that seed dispersal alone cannot explain the value of gene flow in Sparouine. High pollen dispersal distances are known to attenuate spatial clustering and weaken SGS even when seed dispersal remains limited (Sujii et al., 2021; Vekemans & Hardy, 2004). These dynamics may be considered comparatively favorable for maintaining effective gene exchange across space. However, despite these elevated historical dispersal estimates, genetic diversity metrics did not show higher diversity levels in Sparouine relative to the other sites. This apparent decoupling between dispersal distance and standing diversity suggests that long-term diversity patterns are influenced not only by dispersal but also by demographic history and effective population size (Charlesworth, 2009; Hardy et al., 2006).

In contrast, Nouragues, Regina, and Paracou displayed σ values more consistent with previous ecological observations of *D. guianensis*, particularly regarding seed dispersal distances expected under gravity- and wind-mediated dispersal (Jésel, 2005). The σ_s_ values estimated here exceed previously reported mean seed dispersal distances of approximately 30–50 m. Particular attention should be given to Regina, where seed dispersal distance (σ_s_ = 53.1 m) was approximately half that estimated for Sparouine and Paracou. This lower value may indicate more spatially restricted seed-mediated gene flow in Regina. Regina was subjected to selective logging 12 years prior to sampling, and altered canopy structure may influence regeneration patterns and microsite availability (Carneiro et al., 2011; Fredericksen & Mostacedo, 2000; Fredericksen & Pariona, 2002; Hardy et al., 2019; Meer et al., 1998). While the present study was not designed to formally test anthropogenic impacts, the reduced σ_s_ at Regina suggests that historical dispersal dynamics may differ locally. Paracou presents a distinct configuration: despite exhibiting the strongest spatial genetic structure (*Sp* = 0.0293), its historical gene dispersal distance (σ_g_ = 204.4 m) was the highest after Sparouine. The limited pool of parental genotypes within the plot results in a high proportion of related individuals across the sampled area. As illustrated by the COLONY clustering results (Figure 3), most individuals belong to only a few half-sib families, indicating that recruitment is dominated by a restricted number of parental lineages. Under such conditions, even moderate dispersal distances within the plot will not reduce overall relatedness, because individuals share a common genetic background (Born et al., 2008; Vekemans & Hardy, 2004). Strong SGS may arise from limited effective parental diversity rather than extreme dispersal limitation as currently observed in the early life stage (Chung et al., 2003; Ng et al., 2004). Overall, these results demonstrate that SGS patterns in *D. guianensis* reflect the joint legacy of seed dispersal limitation, pollen-mediated connectivity, and site-specific demographic structure.

### 4.4. Contemporary gene dispersal and their drivers

The integration of maternal identification via plastid haplotypes allowed reconstruction of seed dispersal events and confirmed maternal origin in over 21% of offspring assigned to a parent pair. It should be noted that the percentage of offsprings attributed in Sparouine is lower due to the large dispersal distances of this plot, potentially resulting in a greater number of kinship links from outside the sampling area. Parentage analyses across the four study sites revealed variation in seed and pollen dispersal patterns. Despite the short average, seed dispersal distances occasionally exceeded 300–400 m, indicating rare long-distance events. However, seed and pollen dispersal distances beyond 600 m were not detected, likely due to spatial constraints imposed by the site sizes. Pollination distances up to 0.5–14 km have been reported for other canopy tree species, suggesting that the actual breeding range of *D. guianensis* may extend further than our estimates suggest (Dick et al., 2008).

Seed dispersal was predominantly short-range, with most seeds dispersed within 50 m of the maternal tree. This pattern is consistent with previous ecological observations on *D. guianensis*, which describe the species as primarily dispersed by gravity (P. Forget, 1988; Jésel, 2005). Secondary dispersal by rodents has already been suspected but has never really been clearly demonstrated (P. Forget, 1988). The rare long-distance seed dispersal events observed (> 100m) here cannot be fully explained by rodent-mediated dispersal: agoutis and other caviomorph rodents typically move seeds only a few meters from feeding sites (P.-M. Forget, 1990). Alternative vectors such as birds and parrots may play a more significant role in facilitating longer-distance seed movement. Parrots remove the seeds from the pods while still attached to the tree (Pers.Obs. N. Tysklind, (Jésel, 2005), and birds have been shown to transport seeds over larger and more variable distances (Dick et al., 2008; Gelmi-Candusso et al., 2017). Strong winds could also explain such long-distance dispersal, as is the case for other Fabaceae species (Angbonda et al., 2021; Hardy et al., 2019), but this does not adequately explain the variability in distances and direction. Moreover, the morphology of its pods does not appear strongly adapted to sustained long-distance wind transport (Falcão et al., 2022). The occasional secondary dispersal by animals are supported by nuclear *Sp* values below the average *Sp* observed in tropical trees dispersed by wind, gravity, or scatter-hoarding rodents (mean Sp=0.023, N=9)(Dick et al., 2008; Hardy et al., 2006; F. A. Jones & Hubbell, 2006).

Seed dispersal distances differed among sites, with the longest estimates observed in Sparouine and Nouragues. In contrast, shorter distances were detected in Paracou and Regina. These two sites are more accessible and have experienced different levels of human influence in recent decades (logging and hunting). Changes in vertebrate communities associated with increased accessibility and hunting have been documented in French Guiana (Richard-Hansen et al., 2019; Thoisy et al., 2005; Voss et al., 2001), and altered animal assemblages could potentially influence long distance seed dispersal patterns in *D. guianensis* in case of confirmed secondary dispersal by animals. However, given the limited number of sites and the absence of replicated disturbance categories, these interpretations remain exploratory.

These contrasts may also reflect differences in local ecological context, including variation in landscape configuration, or regeneration dynamics. Differences dispersal dynamics between Paracou and Sparouine may also reflect distinct stages in the demographic trajectory of *D. guianensis* populations (Guitet et al., 2014). In Paracou, aggregation and high reproductive skew suggest that the population could be in an early phase of recolonization, where recruitment primarily occurs near parent trees, leading to strong spatial genetic structure and low within-site connectivity. Sparouine represents a more advanced stage of demographic expansion, having reached a stable plateau. This would explain both the greater dispersal distances observed, and the higher diversity of reproductive clusters present at this site.

### 4.5. Reproductive success and individual contribution to gene flow

Current patterns of reproductive success and individual contributions to gene flow provide valuable insights into the actual reproductive ecology and demographic health of *D. guianensis*. Reproductive output was positively correlated with tree size (DBH) in most populations (Regina, Nouragues, and Paracou), suggesting that larger individuals contribute disproportionately to the next generation as observed in other tropical tree species (Angbonda et al., 2021). Across all populations, maternal and paternal roles appeared broadly balanced, with no significant difference in the number of offspring assigned to each role. This reflects a predominantly outcrossing mating system and an equitable reproductive contribution of both sexes.

The *Gini* coefficient revealed strong reproductive dominance in Paracou and Regina; few individuals contributed disproportionately to the gene pool, leading to large full-sib family clusters, with one or two clusters comprising over 190 to 270 offspring. This reproductive skew reduces the effective number of breeders and may limit genetic diversity of subsequent generations (Ismail & Kokko, 2020). In contrast, Sparouine exhibited a more balanced reproductive output, consistent with greater seed and pollen dispersal and reduced kin clustering. High reproductive skew coincided with shorter contemporary gene dispersal distances and stronger spatial genetic structure, suggesting a feedback loop in which restricted dispersal promotes kin aggregation and reinforces dominance by a few individuals (Angbonda et al., 2021; Widiyatno et al., 2017).

Regina in particular displayed pronounced reproductive inequality, large sibship clusters, and comparatively low seed and pollen dispersal distances. These patterns suggest that the contemporary reproductive dynamics in this plot may be less favorable for maintaining broad genetic contribution among adults. Although overall genetic diversity remains stable, the combination of reproductive skew and spatial clustering could reduce the effective population size if maintained over successive generations. Regina experienced selective logging 12 years prior to sampling, the contemporary reproductive patterns observed may reflect local demographic conditions that could potentially be amplified by past disturbance. In French Guiana, forest management operates on a 65-year cutting cycle (PRFB, 2019). If the observed reproductive skew is not reversed before the next harvest, as seen in other forests with short cutting cycles (Widiyatno et al., 2017) it could exacerbate demographic impacts across generations and compromise long-term population resilience.

While the Gini coefficient is used in animal ecology to quantify reproductive skew (Ross et al., 2020, 2023), its application to trees remains rare. Our results highlight its potential to detect emerging demographic imbalances that may not yet be reflected in classical genetic diversity metrics, offering a valuable tool for forest genetic monitoring.

### 4.6. Contrasting demographic trajectories revealed by complementary approaches

The various population genetic and population dynamic approaches used here differ in temporal resolution: SGS reflects gene dispersal over multiple generations, shaped by long-term demographic stability or even by relatively recent colonisation, as it builds up in just a few generations under restricted dispersal as shown in simulation and empirical studies (Epperson, 2005; Sokal & Wartenberg, 1983; Troupin et al., 2006). Conversely, parentage analyses and reproductive success capture recent, one-generation dispersal events within the sampled size classes.

Across all sites, the combination of these temporal perspectives revealed nuanced demographic contrasts. Paracou exhibited intermediate historical gene dispersal distances but the strongest spatial genetic structure and low genetic diversity. Contemporary analyses further revealed pronounced reproductive skew and relatively short effective dispersal distances, reinforcing the signal of limited effective genetic contribution across individuals. Regina displayed a broadly similar configuration in contemporary dynamics, with strong reproductive inequality and relatively restricted effective gene flow, while historical seed dispersal distances were also among the lowest. These combined signals suggest that Paracou and Regina deserve special attention in long-term monitoring programmes, particularly due to their greater accessibility and potential exposure to anthropogenic pressures. These interpretations must remain grounded in species biology and demographic context, including possible variation in older demographic history and environmental variations (Gargiulo et al., 2024; Kokou, 1994).

In Nouragues, historical SGS metrics were similar to those observed in Paracou and Regina, reflecting the species’ intrinsic tendency toward spatial aggregation under limited seed dispersal (Jésel, 2005). Contemporary metrics indicated more balanced reproductive dynamics. This contrast between the historical structure and the current reproductive configuration could reflect improving demographic dynamics within this reserve, which has been protected since 1995.

Sparouine, by contrast, exhibited consistent signals of high connectivity across temporal scales. It displayed the highest historical gene dispersal distances, relatively weak SGS, high pollen dispersal, and low selfing rates. Contemporary estimates confirmed this pattern, with longer median dispersal distances and more equitable reproductive contributions among adults. These results suggest a distinct demographic history, potentially shaped by broader climatic and landscape processes, as previously reported for western French Guiana populations of *D. guianensis* (Bonnier et al., 2023, 2025; Caron et al., 2000).

This study revealed contrasting demographic trajectories among populations and highlighted the power of combining multiple approaches to assess forest population dynamics across temporal scales, by jointly evaluating long-term dynamic and current reproductive patterns. These results reinforce the importance of monitoring reproductive diversity as an early indicator of effective population size decline, particularly given that genetic erosion in long-lived trees may only become detectable several generations after disturbance events (Gargiulo et al., 2024). Even though the variations observed between sites and time scale do not show significantly demonstrated differences, the combined use of the methods applied provides complementary information for assessing the conservation status of forest trees. Depending on the temporal scale and methodological framework applied, the inferred state of population dynamic and conservation status can differ. This study therefore highlights the necessity of integrating complementary genetic and demographic approaches to obtain a more comprehensive and realistic assessment of forest population health (Brooks et al., 2009). By jointly evaluating long-term dispersal legacies and present-day reproductive patterns, we gain a deeper understanding of population trajectories that would remain partially obscured if only a single method were employed.

## Acknowledgements

This work has benefited from support of a grant from Investissement d’Avenir grants of the ANR (CEBA: ANR-10-LABX-25-01). This work is part of project REGE-ADAPT of the research program FORESTT and received government funding managed by the Agence Nationale de la Recherche under the France 2030 program, reference ANR-24-PEFO-0006. JB benefited from a doctoral stipend co-awarded by the Université de Guyane and ADEME. We are especially grateful to St-Omer Cazal for being a major help during the sampling and DNA extraction for this project. We also thank the Master’s students in the Tropical Rainforest (FTH) module of 2023 for their help during a week of sampling at the Paracou field station, Mélaine Aubry-Kientz for her help during sampling of the Sparouine site, Enrique Saez-Laguna for sampling in Regina, and Nathan Goulard for sampling at Nouragues. Microsatellite development and genotyping by sequencing were performed at the PGTB (‘La Plateforme Génome Transcriptome de Bordeaux’) with the help of Christophe Boury and Erwan Guichoux.

## Author contributions

Designed research: JB, OB, ST, NT, MH; performed research: JB, VT, NT, MH; performed the sampling: JB, NT; design of markers, genotyping, and bioinformatic processing: OL, EC, ZC, JB ; analysed data: JB; wrote the paper: JB, NT, MH. All authors critically read and approved the submitted version of the paper.

## Data availability

The *Dicorynia guianensis* plastome sequence is available on the ENA under accession number xxx. The SSR and plastid DNA datasets along with ecological metadata are available on Zenodo under doi xxx.

