## Supporting information 1 for "Complementary evidence from historical and contemporary gene dispersal reveals contrasting population dynamics in a tropical tree species"

**Mismatch between historical and contemporary gene dispersal indicates recent demographic disturbance in a tropical tree species**

**Supporting information 1 : SSR makers design and polymorphisms**

This file contains all the additional information concerning the microsatellite marker design and quality tests.

1. **DNA extraction**

Genomic DNA was extracted from 50 mg of silica-dried leaf or cambium tissue following a modified CTAB protocol (Doyle & Doyle, 1990). Each sample was individually ground into a fine powder using a mortar and pestle under liquid nitrogen and put in 2 mL Eppendorf tubes. The powder was mixed with 900 µL of preheated CTAB extraction buffer (65°C), containing 2% CTAB, 1.4 M NaCl, 100 mM Tris-HCl (pH 8), 20 mM EDTA, and 1% PVP-40, to which 0.4% β-mercaptoethanol was freshly added. After vortexing, 2 µL of RNase A (PureLink, Invitrogen) was added to degrade RNA. Samples were incubated for 45 minutes at 65°C with occasional inversion, then cooled to room temperature. DNA purification was conducted through two consecutive extractions with 900 µL chloroform:isoamyl alcohol (24:1), each followed by centrifugation at 20,000 g for 10–15 minutes at 4°C. The aqueous phase was transferred to a new tube, mixed with an equal volume of ice-cold isopropanol (stored at –20°C), and incubated for 1 hour at –20°C to precipitate DNA. DNA pellets were recovered by centrifugation (20,000 g, 15 min, 4°C), washed successively with 70% and 95% ethanol (–20°C), and dried for ~1 hour at room temperature. DNA was resuspended in 60 µL of TE buffer low-EDTA (10 mM Tris-HCl, 0.1 mM EDTA, pH 8.0). DNA concentration and purity were assessed using a NanoDrop 8000 spectrophotometer (Thermo Fisher Scientific, MA, USA). Samples with DNA concentration below 30 ng/µL or A260/280 < 1.8 were re-extracted.

2. **Marker design**

Microsatellite loci were developed from the *Dicorynia guianensis* draft reference genome (Schmitt *et al.*, 2024), using the automated pipeline described by Lepais *et al.* (2020) and implemented at the Bordeaux Genome-Transcriptome Facility (PGTB). These markers were designed to target di-, tri- and tetranucleotide motifs. The development pipeline consisted of several steps: Genome-wide detection of microsatellite loci, identifying all perfect SSR motifs across the *D. guianensis* reference genome. Filtering for unique loci, to avoid redundancy and paralogous amplifications. Primer design using the filtered unique loci, generated ~20,000 potential SSR markers. From this initial set, two balanced subsets of 60 markers each were constructed (for eventual multiplex amplification after initial tests), with 50% dinucleotide and 50% tri- or tetranucleotide motifs (Table S1.1). Additional selection criteria included: elimination of motifs composed exclusively of AT or GC dinucleotide repeats (due to potential stutter or low polymorphism), retention of perfect repeats only (no compound or interrupted motifs) and minimum of 10 repeat units per motif, to ensure polymorphic potential. All candidate loci were first tested in simplex PCR on a DNA pool including equal representation from each of the four forest plots. Markers were assessed on 1% agarose gel for amplification success and expected fragment length.

To assess seed-mediated gene flow, we developed a complementary set of chloroplast markers, expected to be maternally inherited in angiosperms like *D. guianensis*. The chloroplast genome was reconstructed using PacBio HiFi long reads (data from Schmitt *et al.*, 2024) and assembled with NOVOPlasty v4.3 (Dierckxsens *et al.*, 2017). The resulting circular contig was annotated via BLAST against the NCBI database, revealing 95% identity with the plastome of *Zenia insignis* (Fabaceae, GenBank: PP882779.1), a species belonging to the Dialioideae subfamily, like *Dicorynia guianensis*. The *D. guianensis* plastome displayed few di- or trinucleotide microsatellites. We therefore used two complementary strategies to identify polymorphic loci: 1. Detection of di- and trinucleotide SSRs using QDD v3.1 (Meglécz *et al.*, 2014) to identify perfect SSR motifs. Although QDD does not detect mononucleotide repeats, it identified a small number of di- and trinucleotide SSRs, from which eight loci were retained as candidate markers for primer design. 2. Mononucleotide SSRs were manually identified by visual inspection of the plastid genome using IVG (Robinson *et al.*, 2011). These approaches identified 34 mono, 8 dinucleotide repeats and 7 candidate SNPs for primer design (Table S1.1). All primer pairs were tested in simplex PCR on a DNA pool with equal representation from the four forest plots. Markers were evaluated on agarose gel for amplification success and expected fragment length.

3. **Genotyping**

Genotyping was performed at the Bordeaux Genome-Transcriptome Facility (PGTB) using a high-throughput SSRseq protocol based on Illumina sequencing. Among the individually tested 120 nuclear SSRs and 49 cpSSRs (Table S1.1), the loci showing consistent amplification, clean profiles, and expected product sizes were retained for multiplexing and sequencing (Tables S1.2, S1.3). This resulted in a set of 69 nuclear SSR and 26 plastid markers (allele sequences are given Tables S1.4, S1.5).

The genotyping method consists of three key stages:

- Multiplex amplification (PCR1): Retained loci were grouped into multiplex primer sets. Separate reactions were prepared for nuclear (two sets) and chloroplast markers (one set). A first round of PCR was conducted using 5x HOT FIREPol® MultiPlex Mix (Solis BioDyne) to amplify the three multiplexes on all samples distributed in 384-well plates, using 10 ng of genomic DNA per sample.
- Indexing and adapter ligation (PCR2): after pooling the amplicons from the three multiplexed PCR for each sample, a second PCR was performed to attach dual Illumina indexes (for sample identification) and sequencing adapters to each product of PCR1. Indexing success and amplification quality were verified by agarose gel electrophoresis on a subset of samples.
- Library purification, pooling, and sequencing: Amplicons were pooled by plate using equimolar volumes and purified using AMPure XP magnetic beads (Beckman Coulter). Quality control was performed via Qubit fluorometry and fragment analysis on TapeStation (Agilent). Final libraries were quantified by qPCR and sequenced on an Illumina iSeq 100 2 × 150 bp, following the standard Illumina protocol.

Amplicon reads were demultiplexed by individual, paired-reads were merged using BBmerge (Bushnell *et al.*, 2017) and allele calling was performed using a dedicated SSRseq bioinformatic pipeline integrating FDStools v.1.2.0 (Hoogenboom *et al.*, 2017) described in Lepais et al. (2020).

4. **Marker and dataset quality and polymorphism**

A total of 69 nuclear markers were genotyped, of which 66 with an allelic error rate of <4% (based on repeated genotyping of 95 individuals) and <25% missing data were retained for downstream population genetic analyses (Table S1.2). This selection ensured the robustness and quality of the data obtained for nuclear genetic variability. A total of 26 plastid loci were successfully genotyped, and 23 loci with <25% missing data were retained for analysis (Table S1.3).

To identify problematic samples, e.g. due to DNA contamination, we applied two complementary quality control analyses on each forest plot. First, we calculated individual multilocus heterozygosity with adegenet (Jombart, 2008), defined as the proportion of heterozygous loci per individual across all nuclear SSRs. Individuals with heterozygosity values exceeding the mean plus two standard deviations were considered as outliers, potentially resulting from DNA contamination. Second, we computed pairwise Nei’s genetic distances using the nei.dist() function from the poppr package (Kamvar *et al.*, 2014), after imputing missing data with the most frequent allele at each locus within the plot. Individuals showing abnormally high genetic similarity, defined as Nei’s distance lower than 0.2, were flagged as potentially redundant, which could result from sample mislabeling, clonality, or cross-contamination. Individuals suspected of contamination were removed from subsequent analyses. A total of seven individuals were removed from Sparouine, two from Paracou, six from Nouragues, and one from the Regina site. The removal of these individuals ensured the reliability of subsequent genetic analyses.

Allelic sequences obtained for the 23 plastid loci were concatenated for each individual to reconstruct chloroplast haplotypes. These haplotypes were used in downstream analyses of maternal lineage diversity and seed-mediated gene flow. For individuals with missing data at one or more loci, imputation was performed by replacing the missing allele with the most frequent allele observed at that locus within the same population. This conservative approach allowed us to minimize haplotype loss while limiting the introduction of artificial variation. Haplotype frequencies are provided in Table S1.6.

**Table caption**

The following tables are presented in the *Supporting_information_1_table.xlsx* file

**Table S1.1. Characteristics of the nuclear and chloroplast microsatellite markers used for genotyping.** This table lists the 169 microsatellite markers developed and tested in this study, including 120 nuclear SSRs and 49 chloroplast SSRs. For each marker, we report its identifier, genomic location (or read ID), repeat motif, expected PCR product size (in base pairs), primer sequences, and the multiplex PCR set in which it was included.

**Table S1.2. Summary statistics for the final set of genotyped nuclear microsatellite markers.** This table summarizes the quality metrics and allelic diversity of each microsatellite locus retained for final analyses. For each marker, we report the number of distinct alleles detected based on sequence (NallelesSequence) and size (NalleleSize), the missing data rate (MissingRate), the number of observed allelic mismatches between replicated genotypes (AllelicMismatches), the number of diploid genotypes compared (DiploidGenotypesCompared), and the resulting allelic error rate (AllelicError). These metrics were used to validate marker reliability and support the selection of 69 nuclear SSRs included in population genetic analyses.

**Table S1.3. Summary statistics for the final set of genotyped plastid markers.** This table summarizes the quality metrics and allelic diversity of each plastid marker retained for final analyses. For each marker, we report the number of distinct alleles detected based on sequence (N alleles Sequence) and size (N allele Size), the missing data rate (Missing Rate), the number of observed allelic mismatches between replicated genotypes (Allelic Mismatches), the number of haploid genotypes compared (Haploid Genotypes Compared), and the resulting allelic error rate (Allelic Error). These metrics were used to validate marker reliability and support the selection of 26 plastid markers genotyped and combined into plastid haplotypes for population genetic analyses.

**Table S1.4. Full allelic sequence information for nuclear microsatellite markers (nSSRs) genotyped by SSR-Seq.** This table describes all sequence-defined alleles detected across the 66 nuclear microsatellite loci retained for genetic analyses. For each allele, we report: the locus name (Locus), the annotated sequence (AlleleSequenceAnnotated), including repeat motifs detected by FDSTools, the allele code (AlleleSeqCode), a numeric identifier used to format the multilocus genotype matrix for compatibility with population genetic software, the number of individuals carrying the allele (OccurrencesAcrossIndivs), the raw nucleotide sequence of the allele (AlleleSequence), and the allele length in base pairs (AlleleLength), allowing comparison with traditional size-based genotyping. Each allele corresponds to a unique amplicon sequence (haplotype), differing by variation in the microsatellite repeat region, SNPs in flanking sequences, or indels. This table provides a transparent view of allelic diversity and supports accurate downstream recoding strategies tailored to the research question.

**Table S1.5. Full allelic sequence information for chloroplast microsatellite markers (cpSSRs) genotyped by SSR-Seq**. This table provides complete sequence-based information on the alleles detected at the 23 plastid loci (cpSSRs) retained after quality filtering. For each allele, the following attributes are reported: the locus name (Locus), the annotated sequence (AlleleSequenceAnnotated), including the detected repeat motifs as identified by FDSTools or manual inspection, the allele code (AlleleSeqCode), a unique identifier used for genotype matrix construction, the number of occurrences across all individuals in the dataset (OccurrencesAcrossIndivs), the raw allele sequence (AlleleSequence), showing full-length nucleotide information, and the allele length in base pairs (AlleleLength). Each cpSSR allele represents a distinct haplotype, potentially differentiated by mutations in the repeat motif, SNPs, or indels within the amplified region. This sequence-based documentation enables accurate characterization of maternal lineages, facilitates detection of intra- and inter-population variation, and helps avoid scoring biases due to homoplastic alleles when using length-based approaches.

**Table S1.6. Table of haplotype frequencies obtained from chloroplastic markers at the four sampling sites and overall.** Haplotype and alleles codes are also provided in this table.
