## Supporting information 2 for "Complementary evidence from historical and contemporary gene dispersal reveals contrasting population dynamics in a tropical tree species"

**Mismatch between historical and contemporary gene dispersal indicates recent demographic disturbance in a tropical tree species**

**Supporting information 2 : Regional genetic structure and genetic diversity**

This file contains additional information on the results of the analyses of the regional genetic structure and genetic diversity observed between the 4 study plots presented: Sparouine, Paracou, Nouragues and Sparouine.

**Results from regional genetic structure analysis**

The genetic population structure of *D. guianensis* across the four sampling sites revealed a marked differentiation. In STRUCTURE (Pritchard *et al.*, 2000) analyses at K=2, Paracou and Nouragues were inferred to belong to the same first genetic cluster, while Regina and Sparouine had the highest membership in the second cluster, reflecting a potential genetic separation (Fig. S2.1, S2.2). At K=4, further subdivisions appeared, each sampling site featuring a separate cluster. Pairwise *F_ST_* values revealed weak to moderate differentiation, e.g., between Paracou and Nouragues (*F_ST_* = 0.0457, 134 km), or Nouragues and Regina (*F_ST_* = 0.0639, 65 km)(Table S2.1, S2.2). In contrast, the western population of Sparouine displayed the strongest genetic differentiation from other sites, notably from Paracou (*F_ST_* = 0.2202, 137 km) and Nouragues (*F_ST_* = 0.2016, 212 km), which may be attributed to phylogeographic colonisation patterns or strong genetic drift.

|  | **Paracou** | **Nouragues** | **Regina** | **Sparouine** |
| --- | --- | --- | --- | --- |
| **Paracou** | 0 | 134 | 161 | 137 |
| **Nouragues** | 134 | 0 | 65 | 212 |
| **Regina** | 161 | 65 | 0 | 266 |
| **Sparouine** | 137 | 212 | 266 | 0 |

**Table S2.1. Geographic distance (in km) between the four studied forest plots.**

| **ALL LOCI** | **Paracou** | **Nouragues** | **Regina** | **Sparouine** |
| --- | --- | --- | --- | --- |
| **Paracou** |  | 0.0457*** | 0.0856*** | 0.2202*** |
| **Nouragues** | 0.0457*** |  | 0.0639*** | 0.2016*** |
| **Regina** | 0.0856*** | 0.0639*** |  | 0.1307*** |
| **Sparouine** | 0.2202*** | 0.2016*** | 0.1307*** |  |

**Table S2.2. Pairwise *F*_ST_ distances between *Dicorynia guianensis* sampling sites,** * p < 0.05; **p < 0.01; ***p < 0.001, the significance of the values was obtained by performing 10 000 bootstrap replicates.

| **Plot** | **Nb of defined gene copies** | **Nb of haplotypes** | **Effective Nb alleles** | ***H*_E_** |
| --- | --- | --- | --- | --- |
| **All plots** | 1528 | 27 | 4.18 | 0.761 |
| **Regina** | 417 | 10 | 2.25 | 0.556 |
| **Nouragues** | 357 | 13 | 2.21 | 0.548 |
| **Paracou** | 358 | 12 | 2.05 | 0.512 |
| **Sparouine** | 360 | 5 | 2.30 | 0.566 |

**Table S2.3. Genetic diversity of 27 plastid haplotypes for *D. guianensis* populations in four study sites in French Guiana.** Effective Nb alleles, effective number of alleles (Nielsen *et al.*, 2003); expected heterozygosity (corrected for sample size, Nei, 1978).

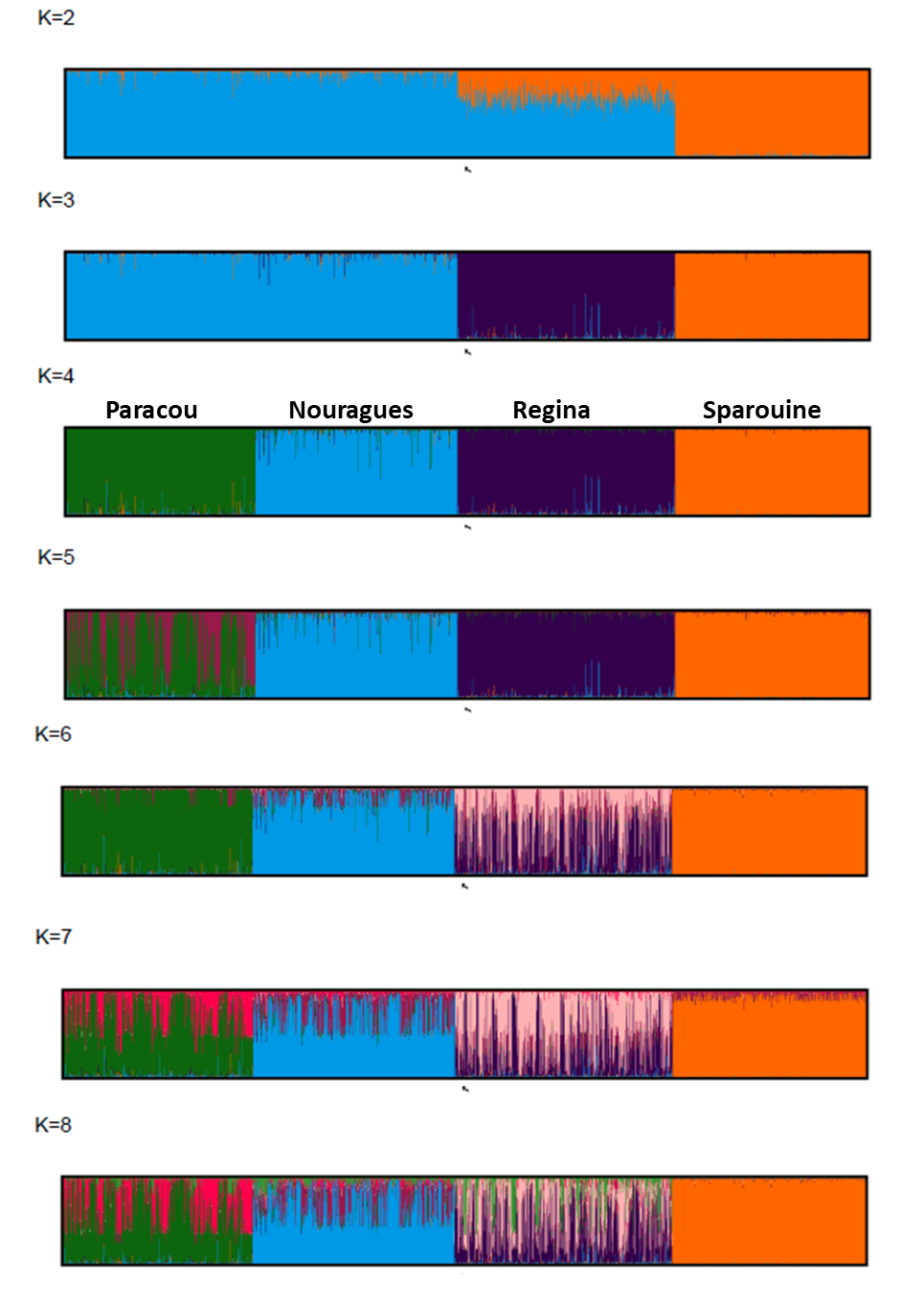

**Figure S2.1. Bayesian estimates of population structure based on nuclear SSR data in sampling sites, from K=2 to K=8.** Each vertical line represents one individual, and color segments represent its assignment proportions to each of K genetic clusters.

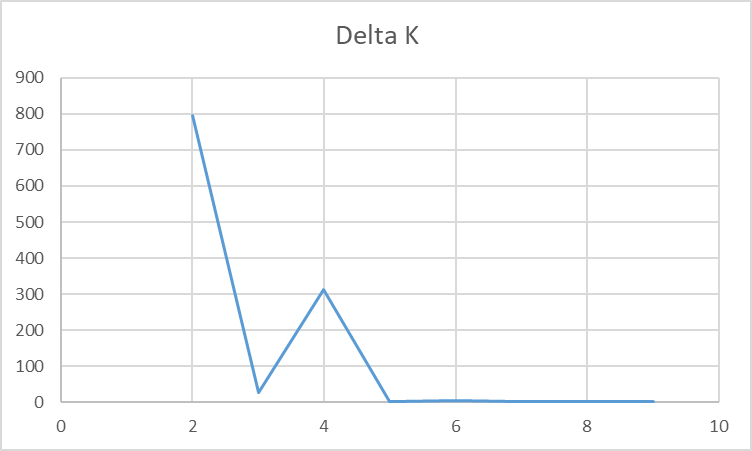

**Figure S2.2. Inference of the optimal number of genetic clusters (K) using the ΔK method on STRUCTURE analyses.** Plot of ΔK values calculated from STRUCTURE runs with K ranging from 1 to 10, based on 20 replicate runs per K, with a burn-in period of 50,000 and 500,000 MCMC iterations. ΔK values were computed using the method of Evanno *et al.* (2005) in Structure Harvester (Earl & vonHoldt, 2012).
