## Supporting information 3 for "Complementary evidence from historical and contemporary gene dispersal reveals contrasting population dynamics in a tropical tree species"

**Mismatch between historical and contemporary gene dispersal indicates recent demographic disturbance in a tropical tree species**

**Supporting information 3 : Fine scale spatial genetic structure and historical gene dispersal**

This file contains additional information on the results of the spatial genetic structure analyses of the four plots studied.

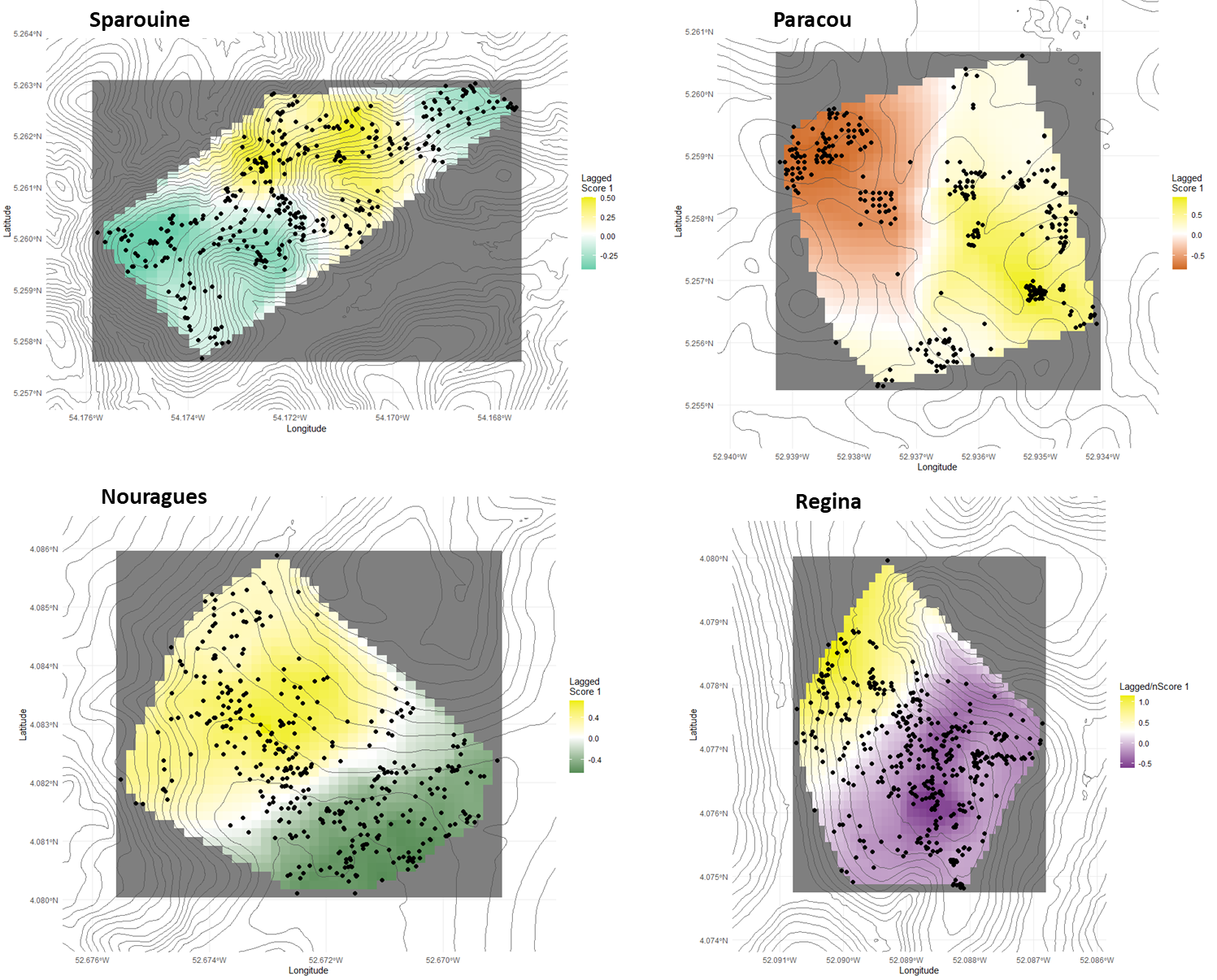

**Figure S3.1. Interpolation map of individual lagged scores of the first global axis of spatial principal component analysis in *Dicorynia guianensis* in the four sampling sites with significant global structures.**

**Figure S3.2. Effect of elevation on spatial genetic structure revealed by sPCA lagged scores in four *Dicorynia guianensis* populations.** Linear regression models were fitted to assess the relationship between individual elevation and their lagged sPCA scores (a proxy for spatial genetic structure) in each study plot. Each panel displays the regression for one plot: Sparouine, Paracou, Nouragues, and Regina. The blue line represents the fitted model. Model statistics (slope, p-value, and adjusted R²) are shown for each site.
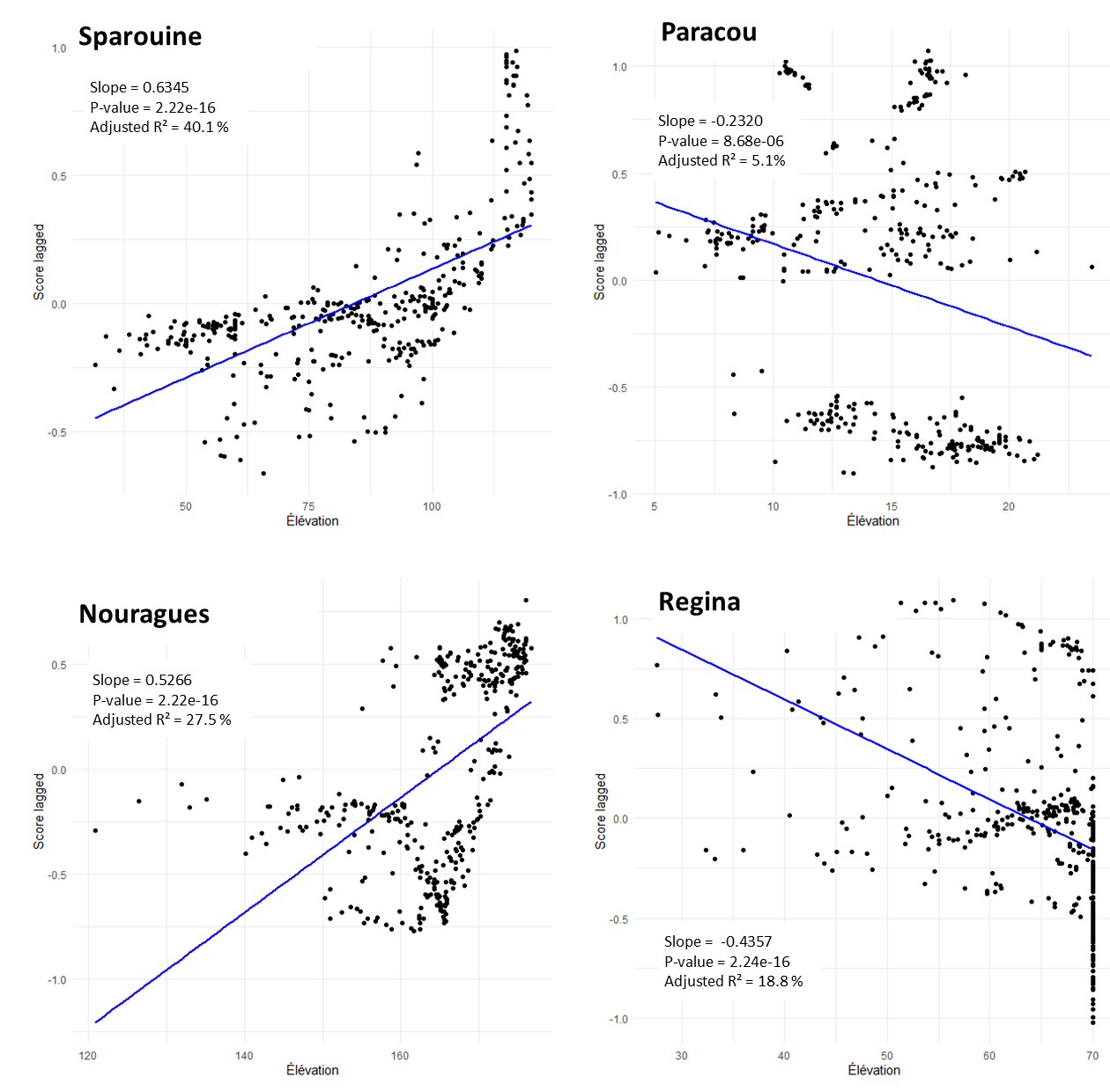

*N.B*

The following figures and tables present results for genetic structure and diversity analyses based on an alternative cohort classification using three size classes: juveniles (JUV, DBH < 4 cm), intermediates (INT, 4 cm ≤ DBH < 30 cm), and adults (ADL, DBH ≥ 30 cm). Although these results are not discussed in the main manuscript, the authors consider them potentially valuable for further reflection on the influence of size class delimitation on the interpretation of cohort-based genetic analyses.

| **Site** | **Size class** | **N** | **Ind/ha** | ***A_R_*** | ***H_E_*** | ***H_O_*** | ***F_IS_*** | **Fi (1st class)** | **b-log** | ***Sp*** |
| --- | --- | --- | --- | --- | --- | --- | --- | --- | --- | --- |
| Sparouine | Juveniles (JUV) | 158 | 5.3 | 4.63 | 0.548 | 0.542 | 0.011 | 0.01 | -0.004 | 0.0041*** |
|  | Intermediate (INT) | 78 | 2.6 | 4.6 | 0.556 | 0.541 | 0.027** | 0.007 | -0.003 | 0.0033*** |
|  | Adult (ADL) | 124 | 4.1 | 4.73 | 0.552 | 0.54 | 0.021** | 0.005 | -0.002 | 0.0021*** |
| Paracou | Juveniles (JUV) | 167 | 6.7 | 4.5 | 0.506 | 0.502 | 0.009 | 0.038 | -0.013 | 0.0135*** |
|  | Intermediate (INT) | 126 | 5 | 4.3 | 0.507 | 0.51 | -0.006 | 0.022 | -0.008 | 0.0079*** |
|  | Adult (ADL) | 65 | 2.6 | 4.18 | 0.493 | 0.498 | -0.011 | 0.037 | -0.014 | 0.0143*** |
| Nouragues | Juveniles (JUV) | 82 | 2.2 | 4.74 | 0.54 | 0.522 | 0.034*** | 0.028 | -0.011 | 0.0109*** |
|  | Intermediate (INT) | 149 | 3.9 | 4.8 | 0.539 | 0.551 | -0.022*** | 0.015 | -0.006 | 0.0062*** |
|  | Adult (ADL) | 146 | 3.8 | 4.63 | 0.537 | 0.577 | -0.074*** | 0.023 | -0.008 | 0.0086*** |
| Regina | Juveniles (JUV) | 215 | 12.6 | 4.63 | 0.563 | 0.553 | 0.018** | 0.02 | -0.009 | 0.0092*** |
|  | Intermediate (INT) | 67 | 3.9 | 4.96 | 0.568 | 0.564 | 0.007 | 0.01 | -0.005 | 0.0051*** |
|  | Adult (ADL) | 135 | 7.9 | 4.85 | 0.56 | 0.552 | 0.014 | 0.011 | -0.005 | 0.0051*** |

**Table. Genetic diversity and spatial genetic structure parameters for D. guianensis individuals categorized into three size classes: juveniles across four study sites in French Guiana.** N, number of individuals sampled; Ind/ha, density of individuals per hectare; *A*_R_ , allelic richness standardized to 50 gene copies; *H_E_*, expected heterozygosity; *H_O_*, observed heterozygosity; *F_IS_*, inbreeding coefficient; sign, significance of F_IS_ (***, P < 0.001; **, P < 0.01; *, P < 0.05; ns, not significant); Fi (1st class), average pairwise kinship coefficient within the first distance class; b-log, regression slope of Fi on the logarithm of distance; Sp, strength of spatial genetic structure and significance (***, P < 0.001).

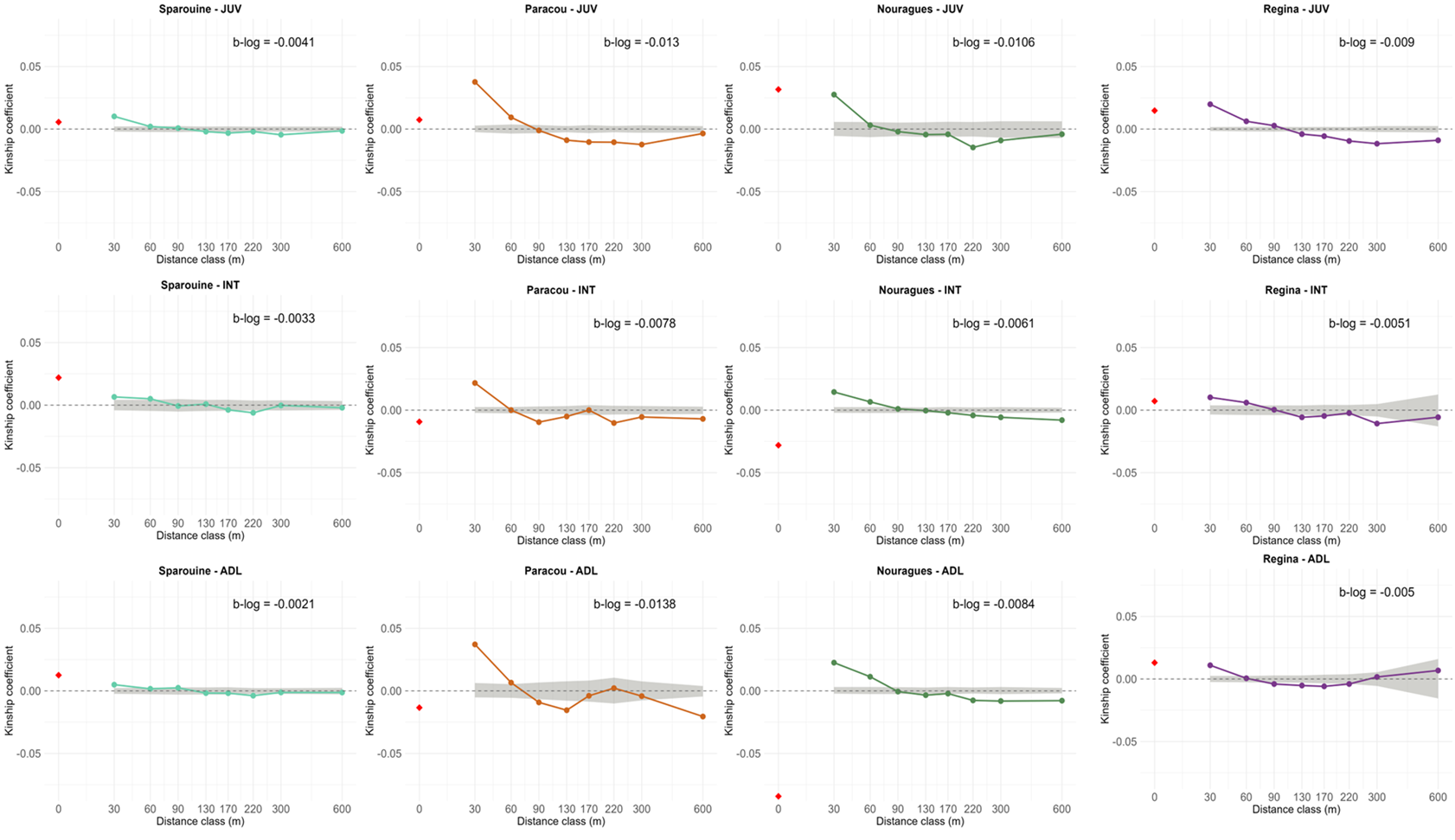

**Figure. Spatial autocorrelograms of the kinship coefficient (Fij) as a function of geographic distance for the four *D. guianensis* study plots for each of the three different cohorts (JUV, INT, ADL).** The dashed grey area represents the 95% confidence interval for spatial randomness of genotypes, generated by 10,000 permutations of individual locations. The regression slope of kinship on log(distance) (b-log) is indicated for each plot. The red rhombus represents the intra-individual kinship coefficient.
