## Supporting information 4 for "Complementary evidence from historical and contemporary gene dispersal reveals contrasting population dynamics in a tropical tree species"

**Mismatch between historical and contemporary gene dispersal indicates recent demographic disturbance in a tropical tree species**

**Supporting information 4 : Parentage analysis and reproductive success**

This file contains additional information on the results of parentage and reproductive success analyses.

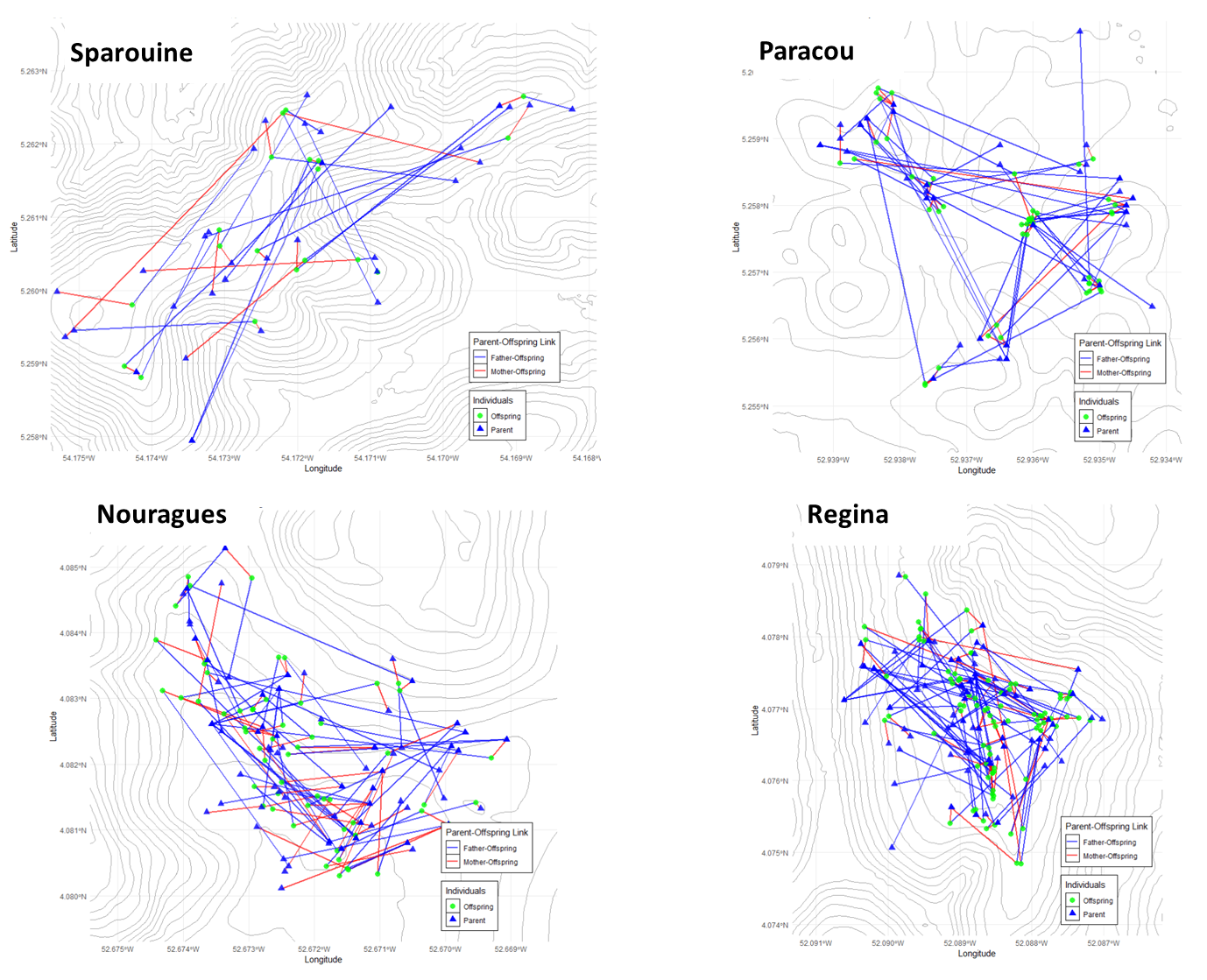
**Figure S4.1. Spatial distribution of parent-offspring relationships inferred from integrated parentage analyses in four *Dicorynia guianensis* forest plots.** Maps showing individual trees (triangles: parents; squares: offspring) and parent-offspring connections for the four study plots (Sparouine, Paracou, Nouragues, Regina). Parentage assignments were derived from concordant results between CERVUS and COLONY software and refined using maternal identity inferred from plastid DNA haplotypes. Blue lines indicate father–offspring links, red lines indicate mother–offspring links (confirmed via plastid haplotype concordance).

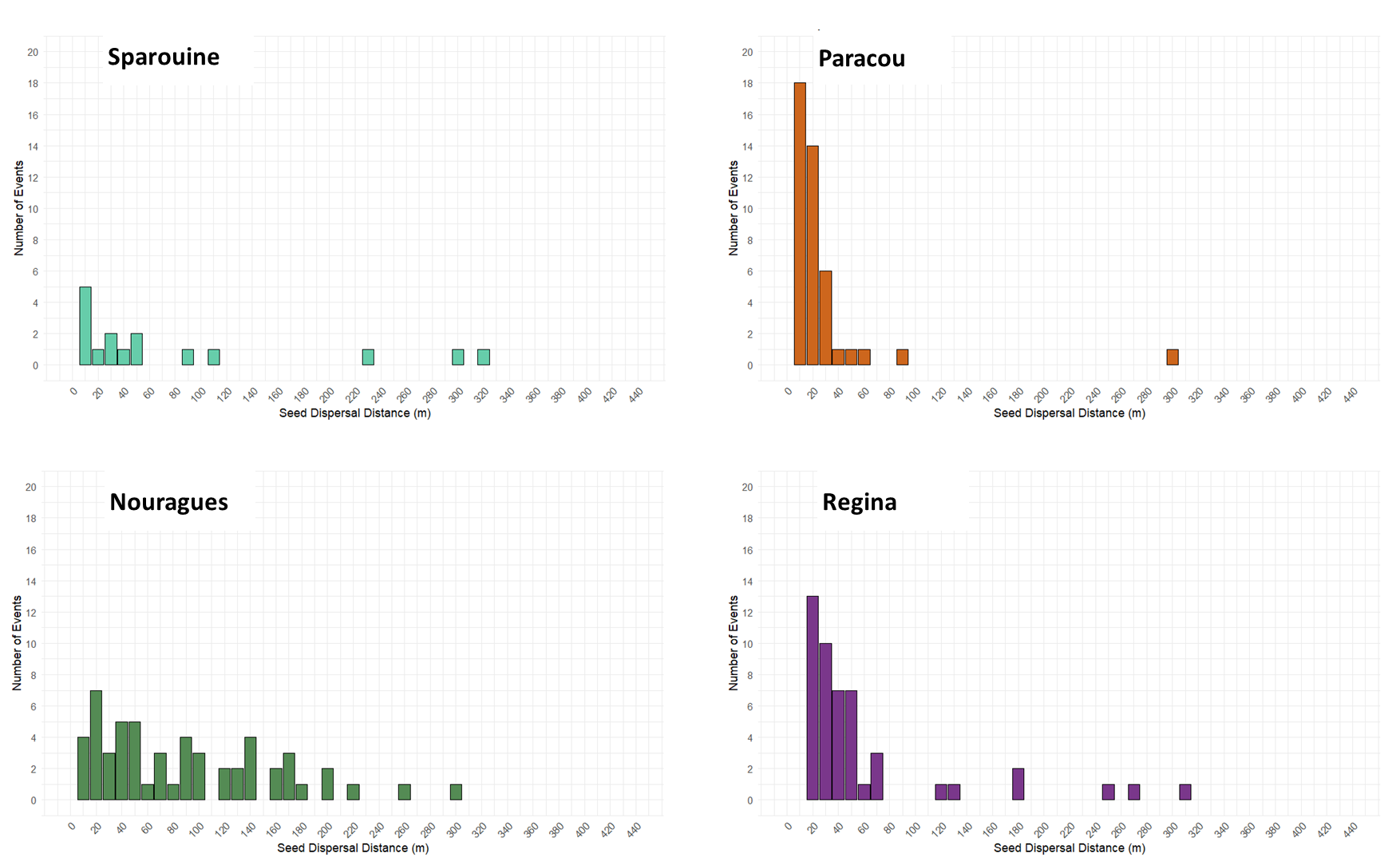
**Figure S4.2. Distribution of seed dispersal distances across four studied plots.** Histograms showing the distribution of seed dispersal distances (in meters) inferred from parentage analysis (COLONY and CERVUS) in each of the four study plots: Sparouine, Paracou, Nouragues, and Regina. Each bar represents the number of inferred seed dispersal events falling within a given distance class (10 m bins). Only offspring with confirmed maternal assignment (based on chloroplast haplotypes) are included.

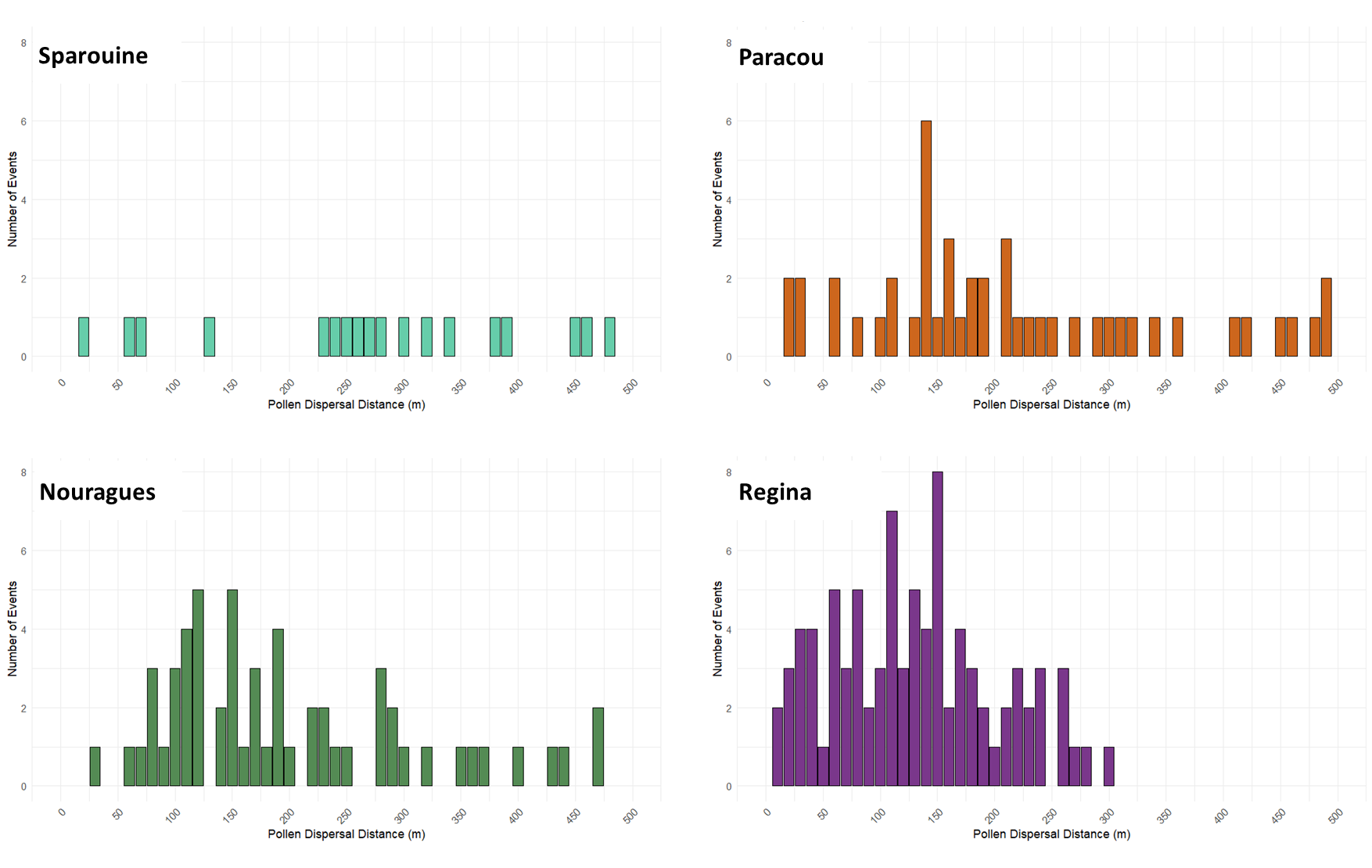
**Figure S4.3. Distribution of pollen dispersal distances across four studied plots.** Histograms showing the distribution of pollen dispersal distances (in meters) inferred from parentage analysis (COLONY and CERVUS) in each of the four study plots: Sparouine, Paracou, Nouragues, and Regina. Each bar represents the number of inferred pollen dispersal events falling within a given distance class (10 m bins). Only offspring with confirmed maternal assignment (based on chloroplast haplotypes) are included.

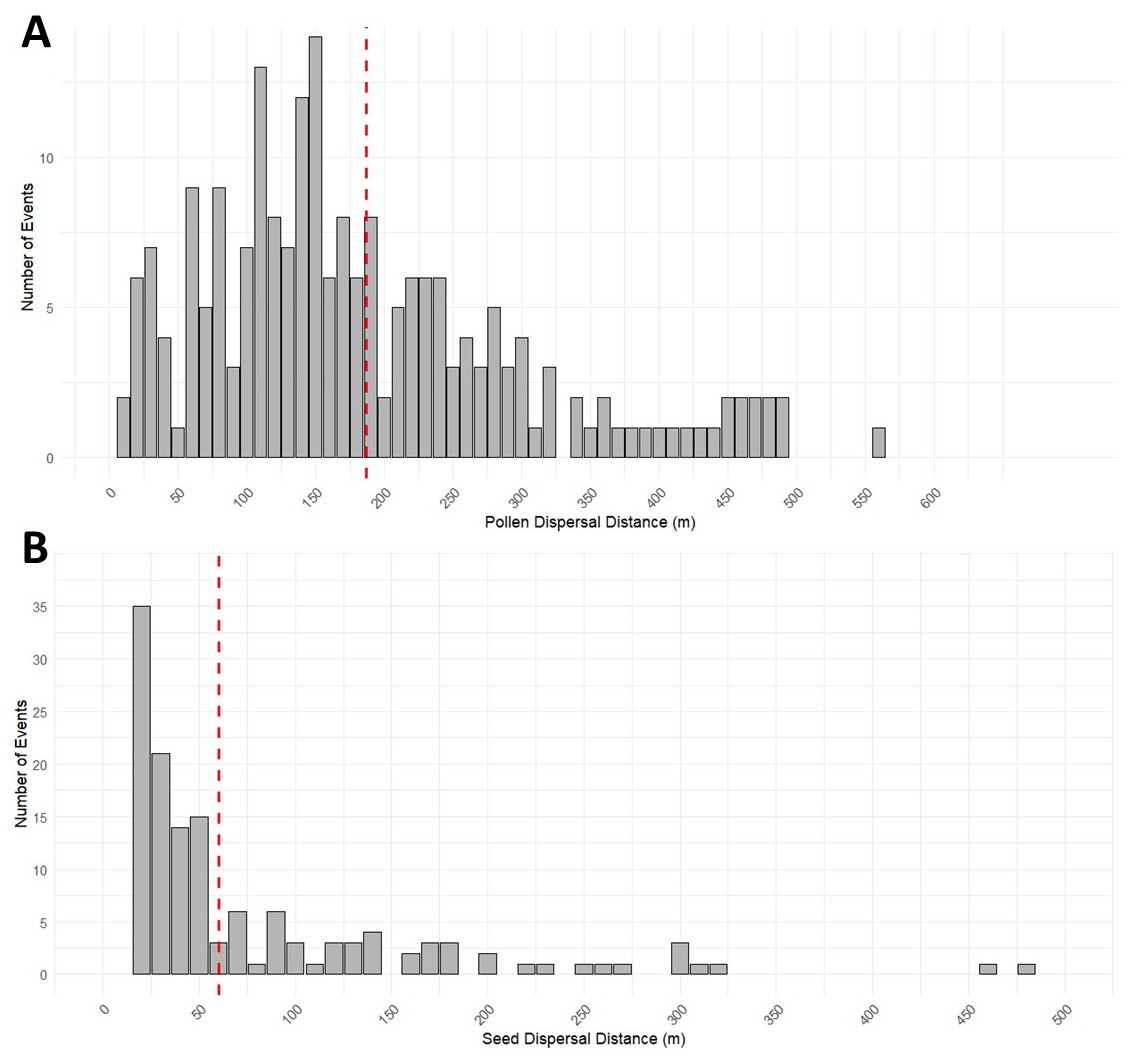

**Figure S4.4.** Overall distribution of pollen and seed dispersal distances. **A.** Barchart showing the distribution of pollen dispersal distances (in meters) across all four study plots combined. The red dashed line represents the mean pollen dispersal distance (205 m). **B.** Barchart showing the distribution of seed dispersal distances (in meters) across all four study plots combined. The red dashed line indicates the mean seed dispersal distance (62 m).

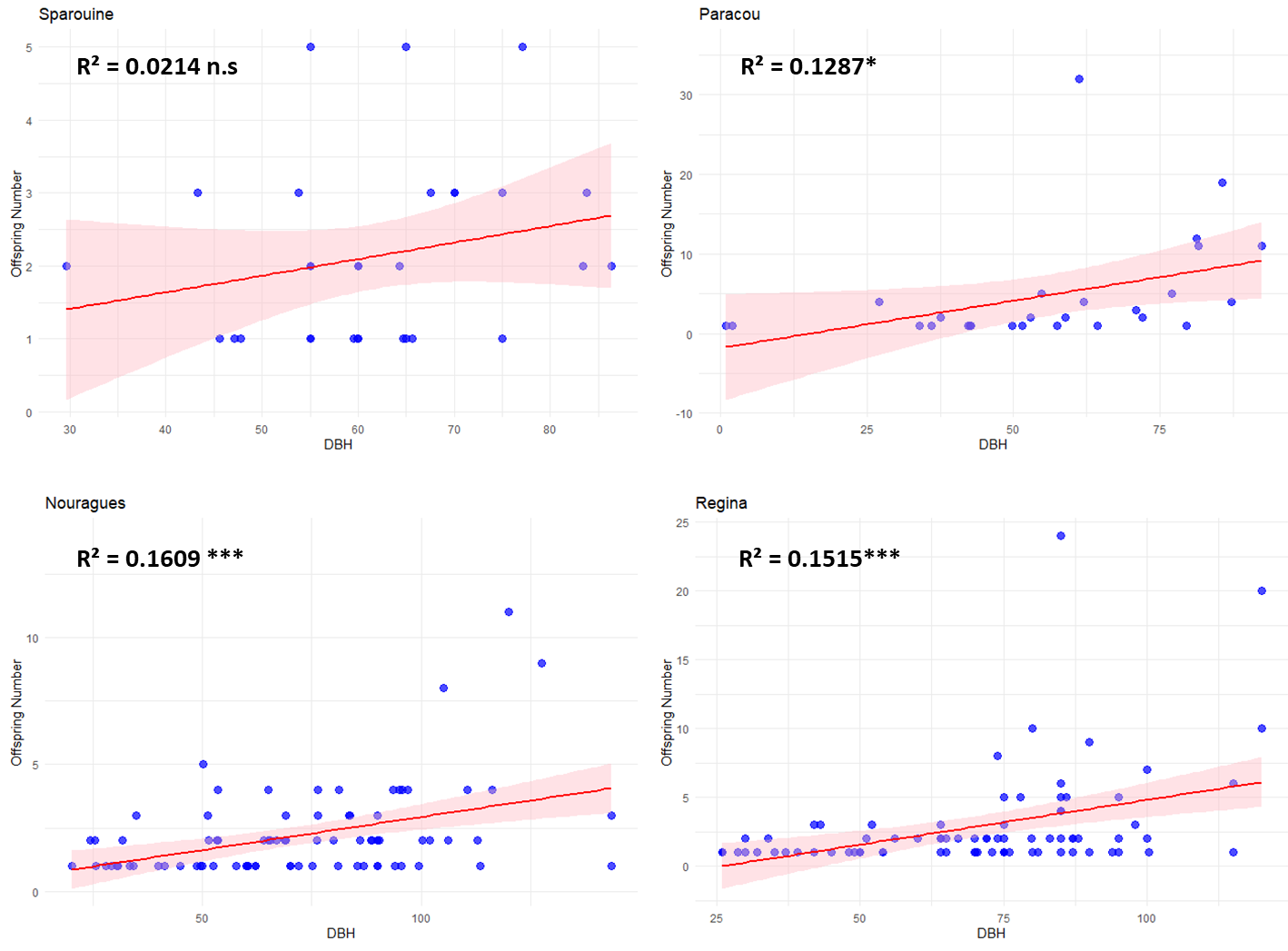

**Figure S4.5. Linear regression analysis between DBH and reproductive success of individuals with DBH over 30cm for the four sampling plots.** The R² values are displayed on each graph in bold, significance of R² value : *, P<0.05; **, P<0.01; ***, P<0.001.

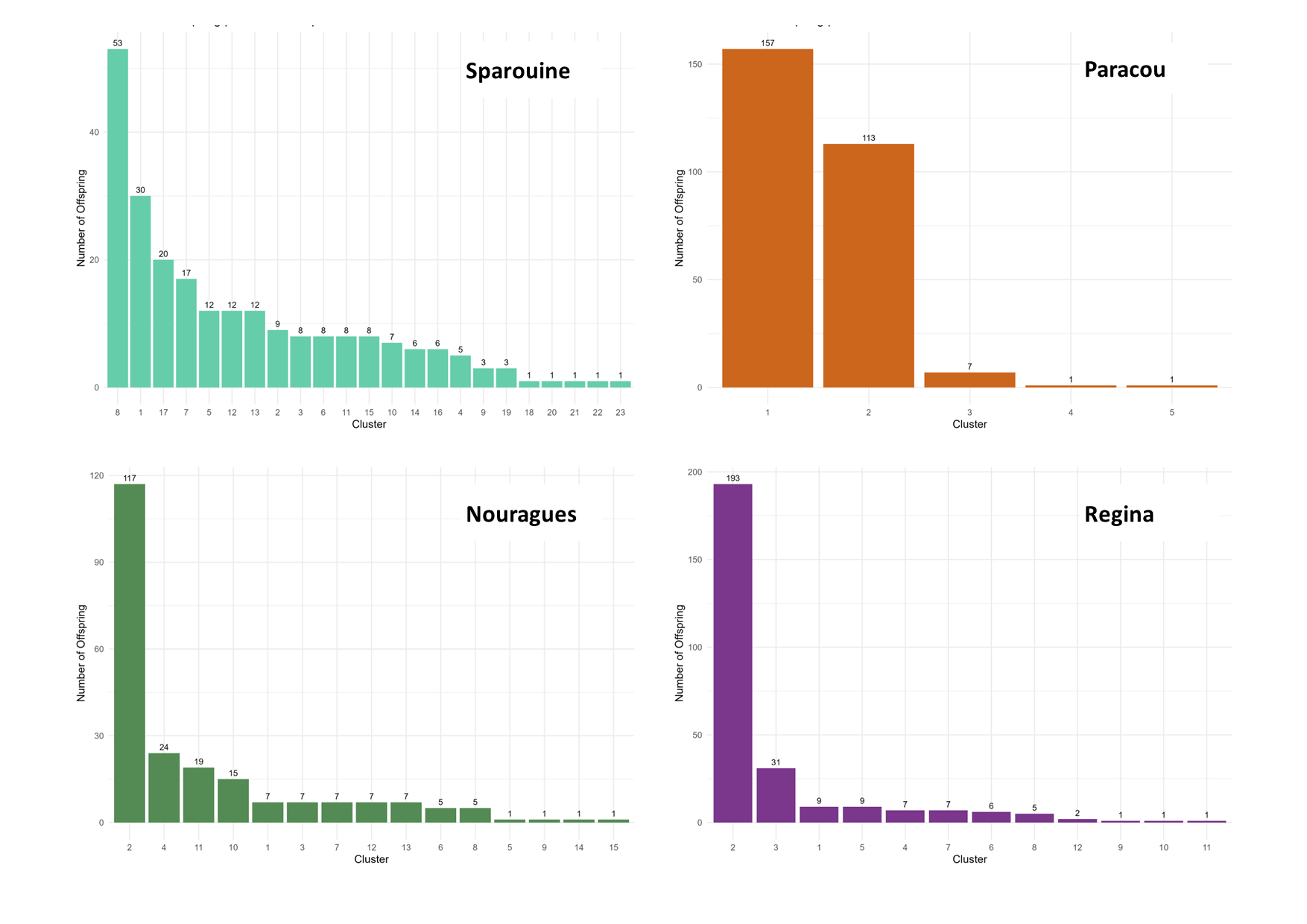
**Figure S4.6. Reproductive cluster sizes across four forest plots.** Barplots showing the number of offspring per reproductive cluster as inferred by COLONY for the four study sites: Sparouine, Paracou, Nouragues, and Regina. Each bar represents a distinct cluster, and its height corresponds to the total number of offspring assigned to that cluster.
